# Oligodendrocyte ER-phagy Restricts Excessive Myelination by Degrading Sphingolipid Synthases

**DOI:** 10.64898/2026.10.04.756522

**Authors:** Tingxin Zhang, Daniela Barbosa, Aksheev Bhambri, Lu O. Sun

**Affiliations:** Department of Molecular Biology, University of Texas Southwestern Medical Center, Dallas, TX 75390, USA; Hamon Center for Regenerative Science and Medicine, University of Texas Southwestern Medical Center, Dallas, TX 75390, USA; Peter O’Donnell Jr. Brain Institute, University of Texas Southwestern Medical Center, Dallas, TX 75390, USA

## Abstract

Myelination is critical for nervous system function. Numerous signaling pathways have been identified that promote myelin sheath growth, disruption of which can lead to hypomyelination, characterized by a reduction in myelin sheath thickness. However, the negative regulators of myelin sheath expansion remain poorly defined. Here, we identify a cellular brake on central nervous system myelination that restricts myelin sheath thickness during brain development. This brake is mediated by a selective form of autophagy, known as endoplasmic reticulum (ER)-phagy, which degrades the ER and its resident sphingolipid synthases. ER-phagy is highly active in oligodendrocytes during myelin membrane expansion, and its impairment leads to ER accumulation, elevated sphingolipid production, and excessive myelin thickening. Genetic reduction of *de novo* sphingolipid biosynthesis in the ER rescues the hypermyelination phenotype caused by ER-phagy disruption. In summary, our findings establish ER-phagy as a key regulator of sphingolipid metabolism and reveal a mechanistic link between ER homeostasis and myelin production by oligodendrocytes, providing insight into mechanisms underlying pathological hypermyelination.

**TEASER:** Oligodendrocyte ER-phagy restrains excessive myelin growth by limiting sphingolipid biogenesis during brain development.

## INTRODUCTION

In the central nervous system (CNS), oligodendrocytes produce one of the largest surface areas of plasma membrane among all cell types, assembling it into the multilamellar myelin sheath (*1*). Due to their large surface area, myelin sheaths comprise nearly 50% of the total dry mass of the human brain (*2*). Myelin sheaths provide critical insulation and trophic support for axons, thereby ensuring proper nervous system function (*3*). The extensive area of myelin sheaths also makes them particularly vulnerable to environmental toxins and activated immune cells (*4, 5*). Therefore, myelin sheath abnormalities are not only found in demyelinating diseases but are also tightly associated with numerous brain disorders and aging (*6*). Despite their significance in health and disease, the precise regulation of myelin sheath growth remains poorly understood.

Numerous signaling pathways have been identified that promote myelin sheath growth, including the phosphoinositide 3-kinase (PI3K)–Akt–mammalian target of rapamycin (mTOR) pathway, the Erk1/2–mitogen-activated protein kinase (MAPK) pathway, neuregulin–ErbB signaling, and cytoskeleton- and neurotransmitter-mediated signaling pathways (*7–19*). Disruption of these pathways can lead to hypomyelination, as evidenced by a reduction in myelin sheath thickness and severe neurological dysfunction. Conversely, hypermyelination has been observed in, and may contribute to, several neurological disorders, such as epilepsy and adult-onset leukoencephalopathy with axonal spheroids (ALSP) (*20, 21*), strongly suggesting that myelin sheath thickness must be tightly regulated to prevent overgrowth. Previous studies have shown that sustained activation of the PI3K/Akt or MAPK signaling pathways in oligodendrocytes significantly increases myelin sheath thickness (*7–13, 16*). Additionally, microglia have been shown to restrict myelin sheath thickness through the TGF-β1–TGF-βR1 axis (*21*). More recently, oligodendrocyte transcription factor EB (Tfeb) was identified as a major repressor of myelination that functions by suppressing cholesterol biosynthesis (*22, 23*). However, the intrinsic cellular and molecular mechanisms that restrict myelin sheath growth remain incompletely understood.

Myelin is rich in lipids, with sphingolipids such as galactosylceramide, sphingomyelin, and sulfatide among its major constituents, contributing to membrane stability, integrity, and compaction (*24*). The establishment of myelination requires the extensive expansion of the myelin membrane, which creates an exceptionally high demand for lipid biosynthesis in oligodendrocytes (*25*). In eukaryotic cells, sphingolipids are predominantly synthesized in the endoplasmic reticulum (ER) and Golgi apparatus (*24*). Meanwhile, the ER, which harbors the largest membrane surface area in the cell, continuously undergoes remodeling through a selective form of autophagy called ER-phagy that enables the targeted degradation of specific ER subdomains (*26*). Despite its important function in maintaining ER homeostasis and regulating neurotransmission in neurons (*27–29*), whether ER-phagy plays a role in myelination remains unknown.

Here, we discover that ER-phagy, an evolutionarily conserved cellular process, is a critical negative regulator of myelin sheath growth. Genetic disruption of autophagy, including ER-phagy, in myelinating oligodendrocytes results in a marked increase in myelin sheath thickness. Mechanistically, impairment of ER-phagy in oligodendrocytes leads to the selective accumulation of ER tubules and ER-resident lipid synthases, thereby enhancing sphingolipid biosynthesis and driving excessive myelin sheath expansion. Conversely, genetic reduction of *de novo* sphingolipid biosynthesis in the ER rescues the hypermyelination phenotype caused by ER-phagy disruption. Together, these findings establish ER-phagy as a key regulator of sphingolipid metabolism in oligodendrocytes and reveal a mechanistic link between ER homeostasis and myelin production.

## RESULT

### Autophagy in myelinating oligodendrocytes limits myelin wrap numbers and sheath thickness

Autophagy is an evolutionarily conserved, lysosomal-mediated process that degrades and recycles cellular components, including proteins, RNA, and organelles such as the ER, thereby preserving cellular homeostasis (Fig. 1A). Previous studies demonstrated that autophagic flux is upregulated during oligodendrocyte differentiation and is critical for oligodendrogenesis and myelin integrity (*30–36*). We asked whether autophagy plays an additional role in regulating myelin sheath growth at the peak of developmental myelination. To genetically disrupt autophagy in premyelinating oligodendrocytes (pre-OLs) and subsequently differentiated myelinating oligodendrocytes, we conditionally deleted Atg5, a molecule essential for autophagosome elongation, using the *Cnp-Cre* driver line (*Cnp-Cre; Atg5^F/F^*; hereafter referred to as *Atg5* cKO; Fig. 1B). We analyzed the ultrastructure of myelin sheaths in the corpus callosum, a major white matter tract where numerous axons are myelinated. At postnatal day 21 (P21), when corpus callosum myelination is at its peak, *Atg5* cKO mice displayed significantly increased myelin sheath thickness compared to littermate controls (*Atg5^F/F^* and *Cnp-Cre; Atg5^F/+^*), as indicated by reduced *g*-ratios (the ratio of inner axonal diameter to the total outer diameter; Fig. 1, C to E and fig. S1, A and B; quantified in Fig. 1, I and K). We observed similar phenotypes in *Cnp-Cre; Atg7^F/F^* mice (hereafter referred to as *Atg7* cKO), in which another autophagosome elongation factor Atg7 was deleted from premyelinating and myelinating oligodendrocytes (Fig. 1, F to H; quantified in Fig. 1, J and K). This increase reflected enhanced myelination rather than decompaction of myelin sheaths at this developmental stage, as both *Atg5* and *Atg7* cKO mice exhibited increased myelin wrap numbers compared to their littermate controls (Fig. 1, L to O). Moreover, we observed no significant differences in the distribution of myelinated axon diameters in *Atg5* and *Atg7* cKO mice compared to controls (fig. S1, C to D’).

**Fig. 1.**
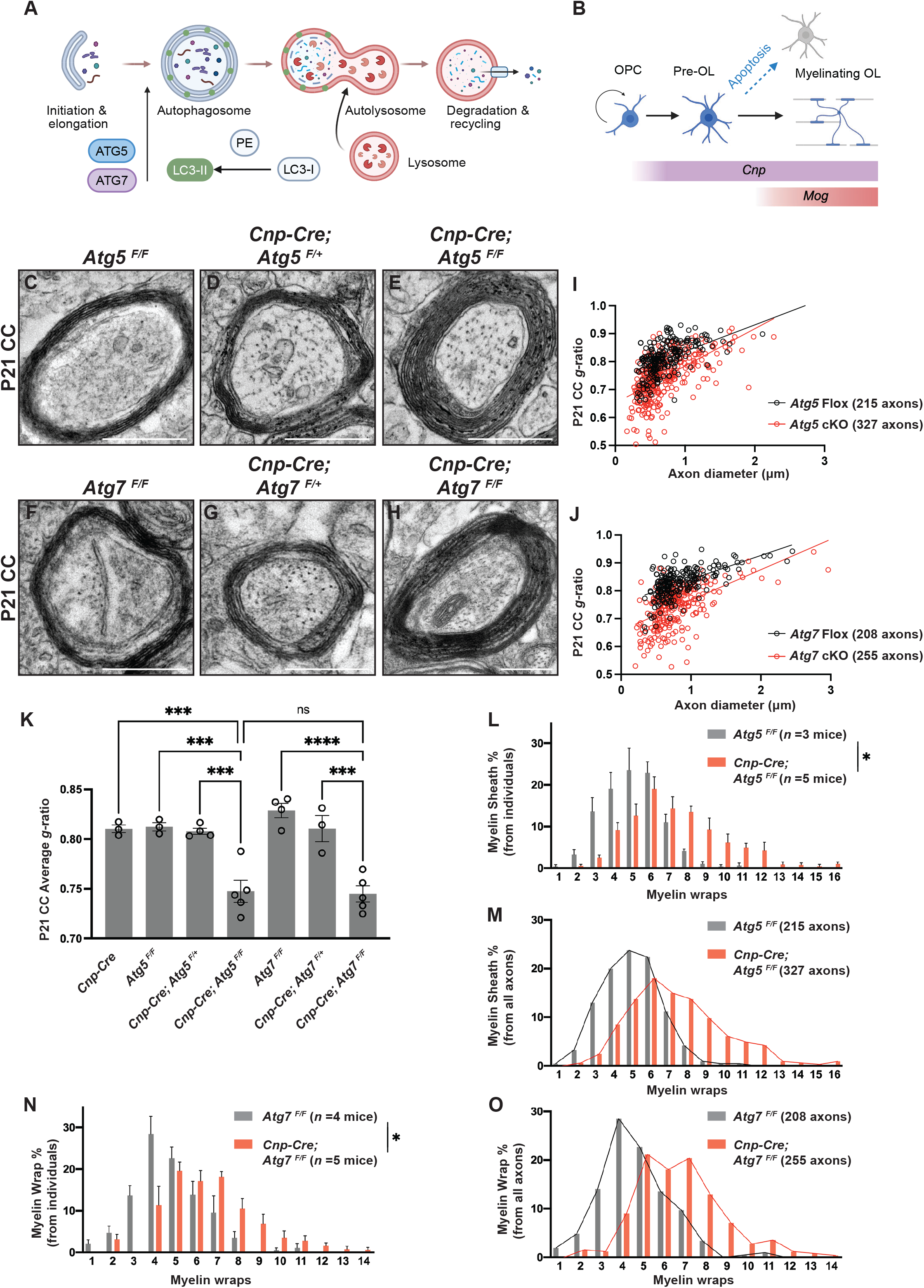
Oligodendrocyte autophagy restricts myelin wrap number and sheath thickness. **(A)** Schematic overview of the autophagy pathway, illustrating autophagosome initiation, elongation, formation, and fusion with lysosomes (adapted from Zhang et al. (*30*); created with BioRender). **(B)** Schematic of oligodendrocyte lineage differentiation from oligodendrocyte precursor cells (OPCs) to premyelinating oligodendrocytes (pre-OLs), and subsequently to mature oligodendrocytes (OLs) (created with BioRender). **(C–H)** Representative transmission electron microscopy (TEM) micrographs from P21 *Atg5^F/F^* (C), *Cnp-Cre; Atg5^F/+^* (D), *Cnp-Cre; Atg5^F/F^* (E), *Atg7^F/F^* (F), *Cnp-Cre; Atg7^F/+^* (G), and *Cnp-Cre; Atg7^F/F^* (H) corpus callosum (CC). **(I and J)** Scatter plot showing *g*-ratios of myelinated axons as a function of axon diameter in P21 *Cnp-Cre; Atg5^F/F^* (*Atg5* cKO) and *Atg5^F/F^*(*Atg5* Flox) mice (I), and the scatter plot showing *g*-ratios from P21 *Cnp-Cre; Atg7^F/F^* (*Atg7* cKO) and *Atg7^F/F^* (*Atg7* Flox) mice (J). **(K)** Quantification of average *g*-ratios of myelinated axons in P21 *Cnp-Cre*, *Atg5^F/F^*, *Cnp-Cre; Atg5^F/+^*, *Cnp-Cre; Atg5^F/F^*, *Atg7^F/F^*, *Cnp-Cre; Atg7^F/+^*, and *Cnp-Cre; Atg7^F/F^* CC. **(L and M)** Quantification of the frequency distribution of myelin wrap numbers from individual mice (L) and from all analyzed axons (M) in P21 *Atg5^F/F^* and *Cnp-Cre; Atg5^F/F^* mice. **(N and O)** Quantification of the frequency distribution of myelin wrap numbers from individual mice (N) and from all analyzed axons (O) in P21 *Atg7^F/F^* and *Cnp-Cre; Atg7^F/F^* mice. One-way ANOVA followed by Tukey’s multiple comparisons test for (K). Kolmogorov-Smirnov test for (L) and (N). Scale bars: 0.5 μm in (C)–(H).

We next investigated whether excessive myelin growth during development generates structural abnormalities observed in aged mice deficient in oligodendrocyte autophagy (*34–36*). We examined established categories of abnormal myelin, including whorls, outfoldings, myelin-on-myelin profiles, and bubbling structures in the corpus callosum (fig. S1E). Quantitative analysis revealed no significant differences in either the total density of abnormal myelin profiles or the density of individual abnormal myelin subtypes in the corpus callosum of *Atg5* cKO mice compared to littermate controls at P21 (fig. S1, F and G), suggesting that the myelin abnormalities observed in adult autophagy-conditional knockout mice are likely due to autophagy’s function in myelin maintenance.

Our previous study showed that both *Atg5* and *Atg7* cKO mice exhibit significantly increased oligodendrocyte numbers due to a reduction in pre-OL apoptosis (*30*). To determine whether an aberrant increase in oligodendrocyte numbers can contribute to myelin sheath thickening, we analyzed *Cnp-Cre; Bax^F/F^; Bak^-/-^* and *Puma^-/-^* mutants, in which pre-OL apoptosis is genetically blocked (*23*). Indeed, P56 *Puma^-/-^* mutants exhibited significantly increased oligodendrocyte numbers in the corpus callosum (fig. S2K). However, neither *Cnp-Cre; Bax^F/F^; Bak^-/-^* nor *Puma^-/-^* mutants displayed alterations in myelin sheath thickness (fig. S2, A to D and G to J), in the distribution of myelin wrap numbers (fig. S2, E and E’) or in the distribution of myelinated axon diameters (fig. S2, F and F’), together showing that aberrantly increasing oligodendrocyte number does not enhance myelin sheath thickness.

To test whether disrupting autophagy in myelinating oligodendrocytes is sufficient to cause aberrant myelin sheath thickening, we leveraged the *Mog-Cre* knock-in line generated by our group, which selectively expresses Cre recombinase in myelinating oligodendrocytes and shows no leakage to other oligodendrocyte lineage stages or other brain cell types (*22*). We generated *Mog-Cre; Atg5^F/F^*mice and characterized myelin sheath thickness and axon diameter distribution in the corpus callosum at P21. *Mog-Cre; Atg5^F/F^* mice displayed significantly increased myelin sheath thickness and a greater number of myelin wraps compared to littermate controls (Fig. 2, A to G and fig. S2, L and M), while the distribution of myelinated axon diameters was comparable (Fig. S2, N and N’). Importantly, unlike *Cnp-Cre; Atg5^F/F^* mutants, which exhibit reduced premyelinating oligodendrocyte apoptosis (*30*), *Mog-Cre; Atg5^F/F^* mutants did not display ectopic myelination in the cerebellar molecular layer or cortex (Fig. 2, H to H’ and J to J’; quantified in Fig. 2, L and N) and showed similar numbers of oligodendrocytes compared to littermate controls (Fig. 2, I to I’ and K to K’; quantified in Fig. 2, M and O). Thus, while autophagy promotes premyelinating oligodendrocyte apoptosis, it has a distinct role in myelinating oligodendrocytes, where it limits myelin wrap numbers and sheath thickness.

**Fig. 2.**
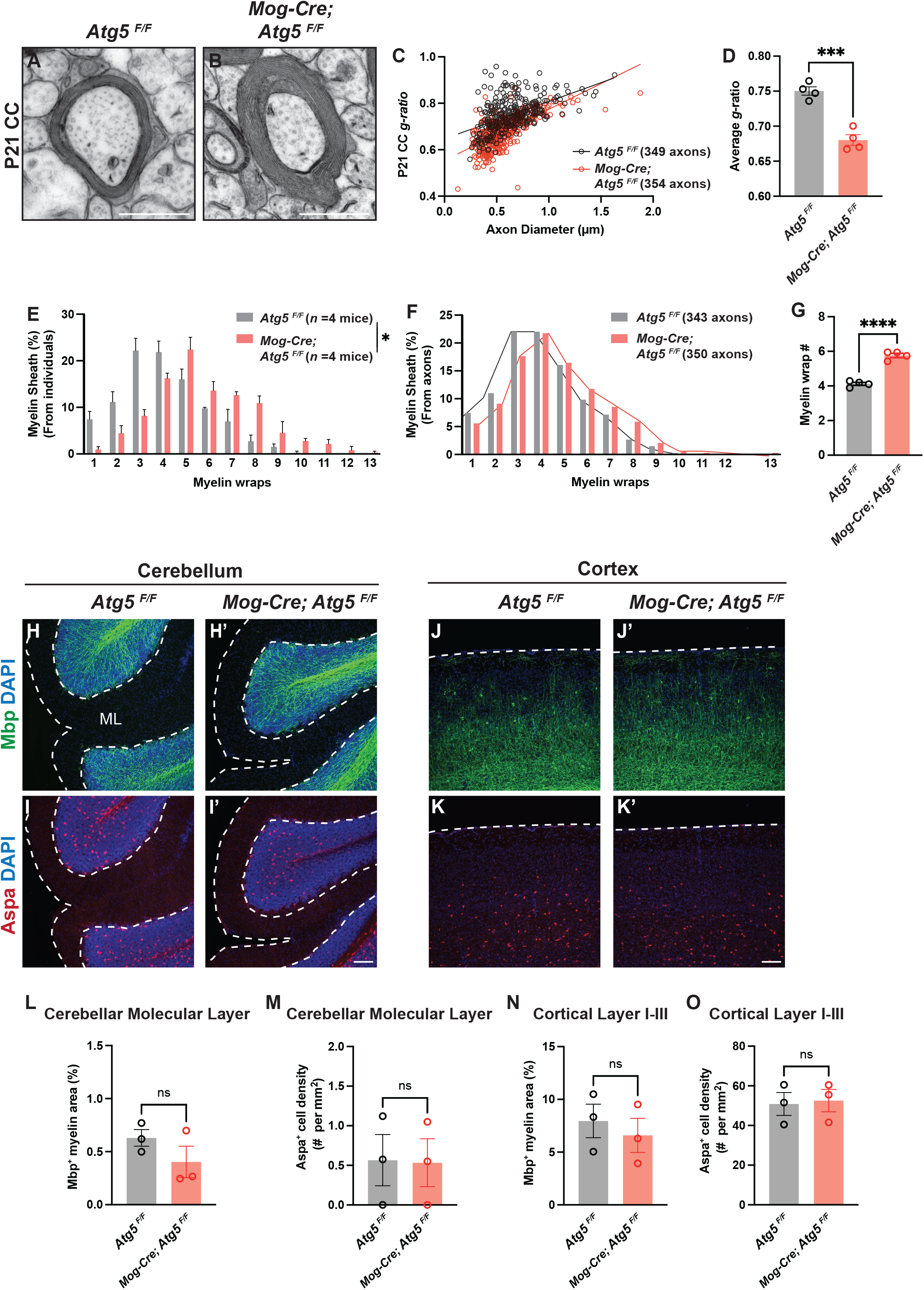
Genetic disruption of autophagy in myelinating oligodendrocytes is sufficient to increase myelin wrap number and sheath thickness without altering oligodendrocyte number. **(A and B)** Representative TEM micrographs from P21 *Atg5^F/F^*(A), *Mog-Cre; Atg5^F/F^*(B) corpus callosum (CC). **(C)** Scatter plot showing *g*-ratios of myelinated axons as a function of axon diameter in P21 *Atg5^F/F^*and *Mog-Cre; Atg5^F/F^* mice. **(D)** Quantification of average *g*-ratios of myelinated axons in P21 *Atg5^F/F^* and *Mog-Cre; Atg5^F/F^* CC. **(E and F)** Quantification of the frequency distribution of myelin wrap numbers from individual mice (E) and from all analyzed axons (F) in P21 *Atg5^F/F^* and *Mog-Cre; Atg5^F/F^*mice. **(G)** Quantification of the average myelin wrap numbers from P21 *Atg5^F/F^* and *Mog-Cre; Atg5^F/F^* CC. **(H*–*I’)** Sagittal brain sections from P21 mice showing myelination in the cerebellar molecular layer (ML), indicated by Mbp fluorescence and Aspa^+^ mature oligodendrocytes in *Atg5^F/F^*mice (H and I) and *Mog-Cre; Atg5^F/F^ mice* (H’ and I’). Dashed lines demarcate the cerebellar molecular layer. **(J–K’)** Sagittal brain sections from *Atg5^F/F^* (J and K) and *Mog-Cre; Atg5^F/F^ mice* (J’ and K’) immunostained by Mbp (green) and Aspa (red) antibodies. Dashed lines indicate the pia surface of the cortex. **(L–O)** Quantification of Mbp^+^ myelin area percentage (L and N) and Aspa^+^ oligodendrocyte numbers (M and O) in the cerebellar molecular layer and cortical layer I–III of P21 *Atg5^F/F^* and *Mog-Cre; Atg5^F/F^* mice. Two-tailed *t*-test for (D), (G), (L), (M), (N), and (O). Kolmogorov-Smirnov test for (E). Scale bars: 0.5 µm in (A) and (B); 100 µm in (I’) for (H)–(I’); and 100 µm in (K’) for (J)–(K’).

### Autophagy selectively degrades ER tubules in oligodendrocytes through ER-phagy during myelin membrane expansion

Autophagy degrades cellular organelles and unwanted proteins to maintain cellular homeostasis (*37*). To investigate the molecular mechanisms underlying autophagy-mediated restriction of myelin sheath thickness, we conducted unbiased quantitative proteomics in *Atg5* KO (*Cnp-Cre; Atg5^F/F^*) and control (*Atg5^F/F^*) oligodendrocytes that had undergone *in vitro* differentiation for 8 days (Fig. 3A). We chose the *Cnp-Cre* driver for mechanistic studies as it allows early and robust gene deletion in the oligodendrocyte lineage (*38*). We focused on the aberrantly enriched proteins in *Atg5* KO oligodendrocytes that would have been degraded by autophagy. By performing gene ontology (GO) analysis of cellular component pathways that were significantly enriched in *Atg5* KO oligodendrocytes, we found that ER-related pathways were among the most highly enriched (Fig. 3B). In addition, among significantly upregulated proteins in *Atg5* KO oligodendrocytes (fold change ≥ 1.5, *p* < 0.05), approximately 35% were ER-resident proteins (Fig. 3C).

**Fig. 3.**
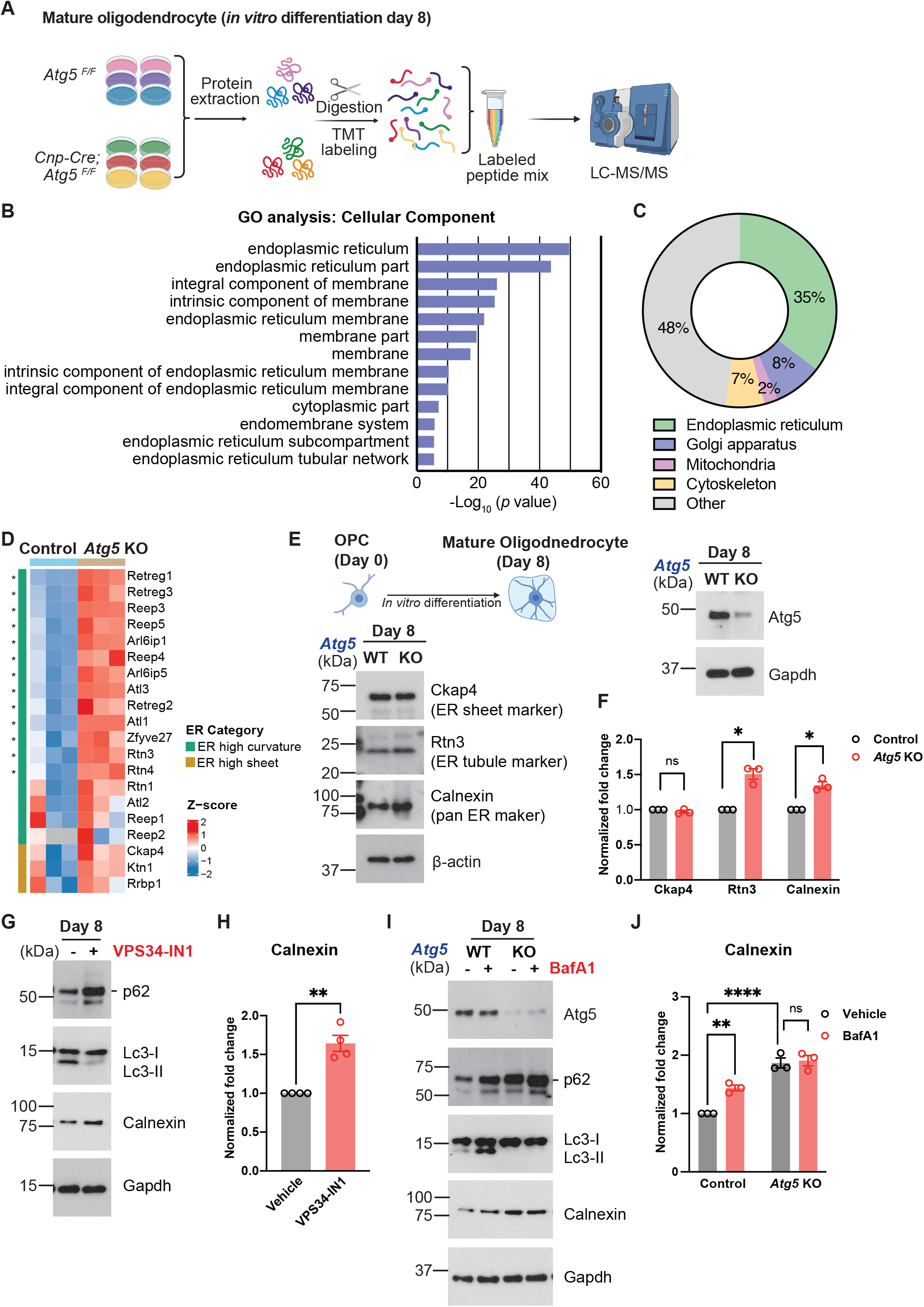
Blockade of autophagy in oligodendrocytes leads to the accumulation of ER tubule–resident proteins. **(A)** Workflow for quantitative proteomics of *Atg5^F/F^* and *Cnp-Cre; Atg5^F/F^* mature oligodendrocytes after 8 days of *in vitro* differentiation (created with BioRender). **(B)** Cellular component gene ontology (GO) analysis of significantly upregulated proteins in the *in vitro* differentiated *Cnp-Cre; Atg5^F/F^* oligodendrocytes versus control (*Atg5^F/F^*) oligodendrocytes at day 8, as revealed by quantitative proteomics. *n* = 3 independent cell purifications. **(C)** Pie chart of significantly upregulated proteins in *Cnp-Cre; Atg5^F/F^* oligodendrocytes, categorized by subcellular component. **(D)** Heatmap of relative protein levels (*z*-scores) for ER high curvature–related proteins (green) and ER sheet–related proteins (brown) in control (*Atg5^F/F^*) and *Atg5* KO (*Cnp-Cre; Atg5^F/F^*) oligodendrocytes, as revealed by quantitative proteomics. Each column represents cells purified from an individual mouse. Each row represents a protein. Significantly increased proteins (*p* < 0.05) are indicated by asterisks. **(E)** Diagram illustrating oligodendrocyte *in vitro* differentiation (top) and Western blots (bottom) showing the protein expression levels of Atg5, Ckap4 (ER sheet marker), Rtn3 (ER tubule marker), and Calnexin (pan-ER marker) in WT (*Atg5^F/F^*) and *Atg5* KO (*Cnp-Cre; Atg5^F/F^*) oligodendrocytes at day 8 *in vitro*. **(F)** Quantification of the normalized fold change of the indicated protein levels in *Atg5* KO oligodendrocytes compared to control oligodendrocytes at day 8 *in vitro*. Protein levels were normalized to β-actin. *n* = 3 independent cell purifications. **(G)** Western blots showing the protein levels of p62, Lc3, and Calnexin in oligodendrocytes with or without VPS34-IN1 treatment. The short line indicates the protein band of p62. **(H)** Quantification of the normalized fold change of Calnexin protein levels in oligodendrocytes treated with VPS34-IN1 compared to those treated with vehicle (DMSO) at day 8 *in vitro*. Protein levels were normalized to Gapdh. *n* = 4 independent cell purifications. **(I)** Western blots showing the protein levels of Atg5, p62, Lc3, and Calnexin in WT and *Atg5* KO oligodendrocytes with or without Bafilomycin A1(BafA1) treatment. The short line indicates the protein band of p62. **(J)** Quantification of the normalized fold change of Calnexin protein levels in oligodendrocytes (WT and *Atg5* KO) treated with BafA1 compared to those treated with vehicle (DMSO) at day 8 *in vitro*. Protein levels were normalized to Gapdh. *n* = 3 independent cell purifications. One-sample *t*-test for (F) and (H). Two-way ANOVA followed by Tukey’s multiple comparisons test for (J).

The ER can be further categorized as ER tubules and sheets based on their shape and function (*39–41*). ER sheets are flat membranes harboring ribosomes, primarily responsible for synthesizing membrane and secreted proteins (*39–41*). By contrast, ER tubules are long, cylindrical structures lacking ribosomes but specialized in lipid synthesis, ion transport, and forming contact sites with other organelles (*39–41*). To determine whether autophagy alters ER proteins unbiasedly or selectively, we analyzed all ER-related proteins that were aberrantly increased in *Atg5* KO oligodendrocytes. ER tubule–associated proteins showed marked upregulation compared to controls (proteins belonging to “ER high curvature” in Fig. 3D; fig. S3), whereas ER sheet–associated proteins exhibited only minor changes in *Atg5* KO oligodendrocytes (proteins belonging to “ER high sheet” in Fig. 3D; see the complete protein list in Table S1). These results were validated using Western blot analysis of the ER sheet marker Ckap4, the ER tubule marker Rtn3, and the pan-ER marker Calnexin (Fig. 3E). Consistent with the proteomic results, Ckap4 levels remained unchanged, while Rtn3 and Calnexin levels were significantly increased in *Atg5* KO oligodendrocytes compared to controls (Fig. 3F).

To determine whether autophagy negatively regulates oligodendrocyte ER abundance *in vivo*, we acutely purified mature oligodendrocytes from mouse brains and detected ER-related proteins using Western blot analysis (fig. S4A). Indeed, the levels of the pan-ER marker Calnexin, as well as the ER-phagy receptors Atl3 and Calcoco1 (*42, 43*), which localize to the ER, were significantly increased in acutely purified mature oligodendrocytes from *Atg5* cKO mice compared with controls (fig. S4, B and C).

To rule out the possibility that the observed ER protein alteration was due to Atg5-dependent but autophagy-independent effects (*44*), we treated oligodendrocytes with two autophagy inhibitors: VPS34-IN1, a selective Vps34 inhibitor that blocks autophagy initiation, and Bafilomycin A1 (BafA1), which prevents autophagosome-lysosome fusion (*45–47*). As expected, VPS34-IN1 treatment reduced the levels of Lc3-II, the lipidated form of Lc3 that marks autophagosomes, while increasing p62 accumulation, a known substrate of autophagy (Fig. 3G). These results indicate effective autophagy inhibition by VPS34-IN1. We found that the ER marker Calnexin levels were significantly increased in oligodendrocytes treated with VPS34-IN1 compared to those treated with the vehicle (Fig. 3G; quantified in Fig. 3H). Similarly, *Atg5* KO oligodendrocytes displayed decreased Lc3-II and increased p62 levels, as well as a lack of further response to BafA1 treatment (Fig. 3I), confirming that genetic deletion of Atg5 in oligodendrocytes disrupts autophagic flux. Importantly, BafA1 treatment significantly increased Calnexin levels in *Atg5* WT oligodendrocytes but not in *Atg5* KO cells (Fig. 3I; quantified in Fig. 3J), together showing that autophagy degrades ER-related proteins in an Atg5-dependent manner.

ER-phagy represents a unique form of autophagy that selectively degrades ER (*48*). To visualize this dynamic process during myelin membrane expansion, we employed a fluorescence reporter assay to monitor ER-phagy flux in live oligodendrocytes (*49*). Oligodendrocytes were infected with a lentivirus expressing the ER-phagy reporter (ss-GFP-RFP-KDEL), which labels the ER with both GFP and RFP signals (Fig. 4A). During autophagic degradation, ER fragments are delivered to lysosomes, where the reporter is cleaved into GFP and RFP fragments. Because GFP fluorescence is rapidly quenched in the acidic environment of lysosomes, while the more stable RFP signal remains detectable, the presence of RFP-only puncta serves as an indicator of ER-phagy events (*49*).

**Fig. 4.**
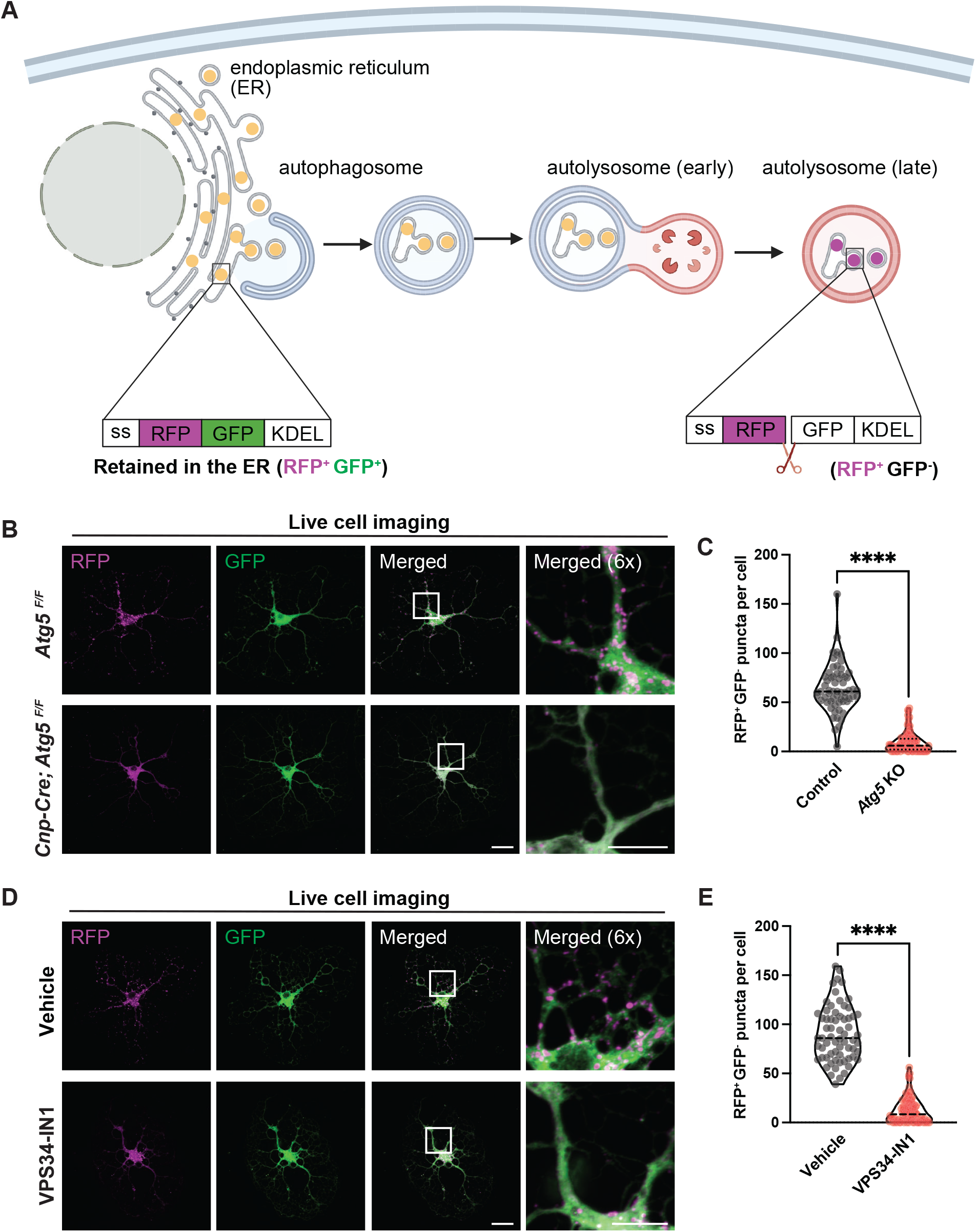
Autophagy mediates ER degradation (ER-phagy) in oligodendrocytes during myelin membrane expansion. **(A)** Schematic of ER-phagy flux detection using the ss-RFP-GFP-KDEL reporter (created with BioRender). ss-RFP-GFP-KDEL is targeted to the ER lumen via an N-terminal ER signal sequence (ss) derived from the ER protein BiP and sequestered by a C-terminal ER-retention peptide (KDEL) (RFP^+^ GFP^+^). During ER-phagy, lysosomal cleavage yields both RFP and GFP fragments. GFP fluorescence is quenched in lysosomes while RFP remains stable, allowing the RFP^+^ GFP^-^ signal to serve as the readout for ER-phagy flux. **(B)** Live-cell imaging of *Atg5^F/F^* and *Cnp-Cre; Atg5^F/F^* oligodendrocytes infected with the ER-phagy reporter (ss-RFP-GFP-KDEL) at day 8 *in vitro.* White insets represent the enlarged views shown on the rightmost column. **(C)** Quantification of RFP^+^ GFP^-^ puncta per cell in *Atg5^F/F^*and *Cnp-Cre; Atg5^F/F^* oligodendrocytes. Each group contains > 60 cells from 3 independent cell purifications. **(D)** Live-cell imaging of oligodendrocytes infected with the ER-phagy sensor followed by VPS34-IN1 or vehicle (DMSO) treatment at day 10 *in vitro*. White insets represent the enlarged views shown on the rightmost column. **(E)** Quantification of RFP^+^ GFP^-^ puncta per cell in oligodendrocytes treated with DMSO vehicle or VPS34-IN1. Each group contains > 60 cells from 3 independent cell purifications. Two-tailed *t*-test for (C) and (E). Scale bars: 10 μm in (B) and (D).

We first evaluated the expression of the ER-phagy reporter in oligodendrocytes following lentiviral transduction. GFP^+^ RFP^+^ double-positive cells expressed the oligodendrocyte marker MBP and exhibited the stereotypical pancake-like morphology at day 8 of differentiation, confirming successful lentiviral transduction in oligodendrocytes (Fig. S5). Next, we subjected the transduced cells to live-cell imaging. We observed numerous RFP puncta in control oligodendrocytes at day 8 of differentiation while they were actively expanding their membrane, indicating active ER-phagy at this stage (Fig. 4B; quantified in Fig. 4C). In stark contrast, *Atg5* KO oligodendrocytes or cells treated with the autophagy inhibitor VPS34-IN1 showed a significant reduction in the number of RFP^+^ GFP^-^ puncta (Fig. 4, B and D; quantified in Fig. 4, C and E). Thus, oligodendrocyte ER-phagy maintains ER homeostasis and reduces ER tubule–associated proteins during myelin expansion.

### Oligodendrocyte ER-phagy limits *de novo* sphingolipid biogenesis to repress myelin sheath growth

ER tubules harbor numerous lipid synthases and are tightly associated with lipid biosynthesis (*50*). To test whether oligodendrocyte ER-phagy degrades ER tubules and associated lipid synthases, we performed GO analysis of the upregulated proteins from *Atg5* KO oligodendrocytes, focusing on biological processes. Indeed, lipid biosynthetic pathways were among the most significantly enriched biological processes (Fig. 5A). Notably, *de novo* sphingolipid synthesis mainly occurs on ER tubules, where ceramides are generated as major precursors for many myelin-enriched lipids such as galactosylceramide, sulfatide, and sphingomyelin (Fig. 5B) (*51*). We found that enzymes involved in sphingolipid synthesis, including Sptlc1/2, Kdsr, Cers2, Degs1, and Cgt, were significantly upregulated in *Atg5* KO oligodendrocytes compared to controls (Fig. 5C). These results were further validated by Western blot analysis using *Atg5* KO oligodendrocytes and VPS34-IN1–mediated pharmacological inhibition of autophagy (Fig. 5, D and E); both Sptlc1 and Cers2 levels were significantly increased in *Atg5* KO oligodendrocytes and in cells treated with VPS34-IN1 compared to controls (Fig. 5, F and G).

**Fig. 5.**
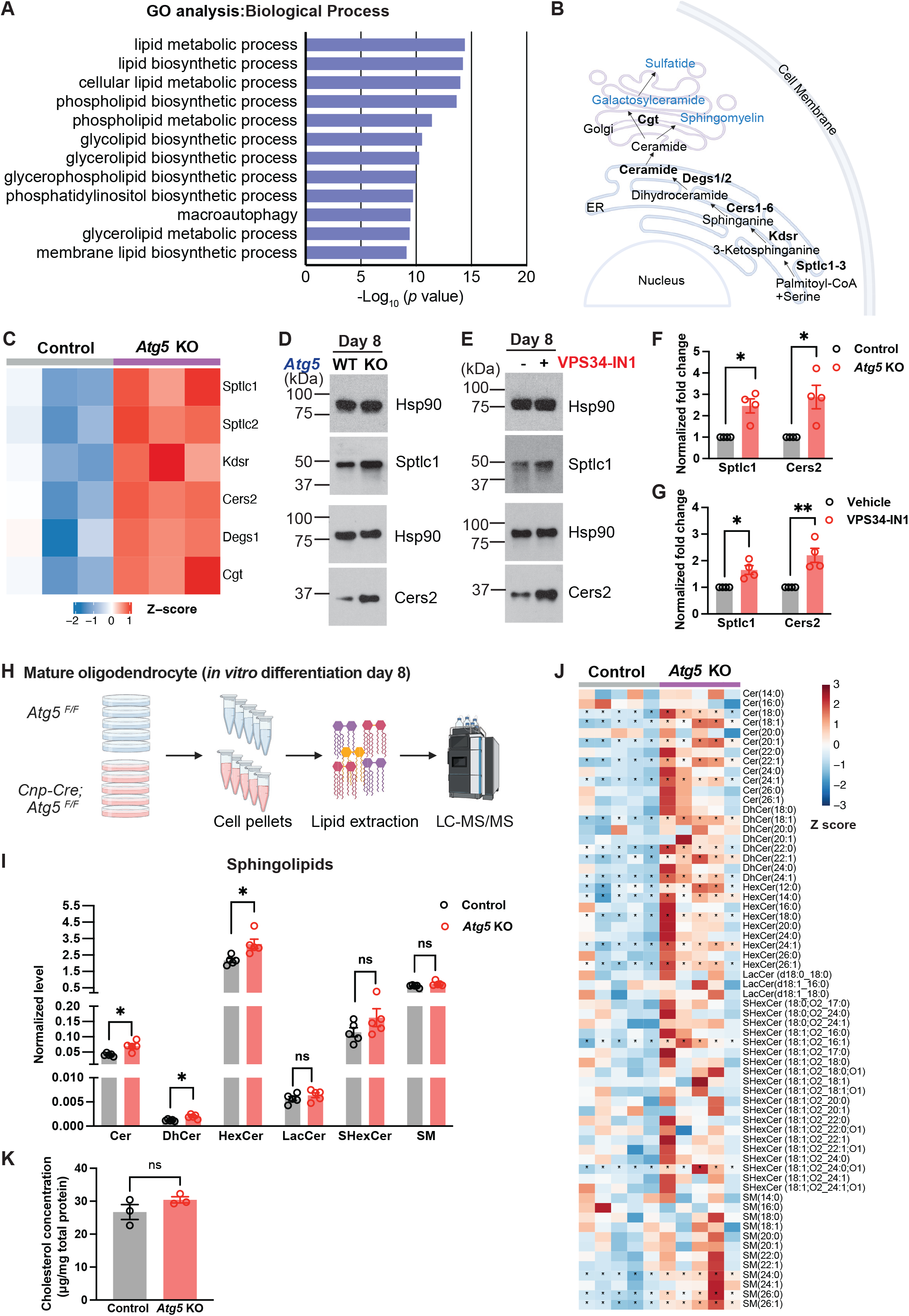
ER-phagy limits *de novo* sphingolipid biogenesis on the ER of oligodendrocytes. **(A)** Gene ontology (GO) analysis of biological processes for proteins significantly upregulated in day 8 *in vitro* differentiated *Cnp-Cre; Atg5^F/F^* oligodendrocytes compared to controls (*Atg5^F/F^*), as revealed by quantitative proteomics. *n* = 3 independent cell purifications. **(B)** Diagram illustrating *de novo* sphingolipid biosynthesis pathways (created with BioRender). **(C)** Heatmap showing relative protein levels (*z*-scores) of *de novo* sphingolipid biosynthetic enzymes from control (*Atg5^F/F^*) and *Atg5* KO (*Cnp-Cre; Atg5^F/F^*) oligodendrocytes at day 8 *in vitro* as revealed by quantitative proteomics. **(D and E)** Western blots showing Sptlc1 and Cers2 protein levels in WT (*Atg5^F/F^*) and *Atg5* KO (*Cnp-Cre; Atg5^F/F^*) oligodendrocytes at day 8 *in vitro* (D), and in oligodendrocytes treated with vehicle or VPS34-IN1 at day 8 *in vitro* (E). **(F and G)** Normalized fold change of indicated protein levels in *Atg5* KO oligodendrocytes compared to WT controls at day 8 *in vitro* (F), and in oligodendrocytes treated with VPS34-IN1 compared to those treated with vehicle at day 8 *in vitro* (G). Protein levels were normalized to Hsp90. *n* = 4 independent cell purifications. **(H)** Workflow of lipidomic analysis for *Atg5^F/F^* and *Cnp-Cre; Atg5^F/F^*mature oligodendrocytes following 8 days of *in vitro* differentiation (created with BioRender). **(I)** Relative abundance of sphingolipid species in control and *Atg5* KO oligodendrocytes, as determined by lipidomics. Normalization was based on lipid reads per total protein level. Cer, ceramide; HexCer, hexosylceramide; LacCer, lactosylceramide; SHexCer, sulfatide; SM, sphingomyelin. *n* = 5 independent cell purifications. **(J)** Heatmap showing relative levels (*z*-scores) of sphingolipid species with varying fatty acid chains from control (*Atg5^F/F^*) and *Atg5* KO (*Cnp-Cre; Atg5^F/F^*) oligodendrocytes at day 8 *in vitro,* as revealed by lipidomics. Each column represents cells purified from an individual mouse. Each row represents a lipid species, distinguished by its fatty acid chain composition. Significantly increased lipids (*p* < 0.05) are indicated by asterisks. **(K)** Quantification of total cholesterol levels in control (*Atg5^F/F^*) and *Atg5* KO (*Cnp-Cre; Atg5^F/F^*) oligodendrocytes at day 8 *in vitro*. One-sample *t*-test for (F) and (G). Two-tailed *t*-test for (I) and (K).

To assess whether aberrantly elevated sphingolipid synthases alter lipid metabolism in *Atg5* KO oligodendrocytes, we performed lipidomic profiling of *Atg5* KO (*Cnp-Cre; Atg5^F/F^*) and control (*Atg5^F/F^*) oligodendrocytes that had differentiated for 8 days *in vitro* (Fig. 5H). As expected, total levels of ceramides, dihydroceramides, and hexosylceramides were significantly elevated in *Atg5* KO cells compared to control cells (Fig. 5I; Table S2). Notably, compared to species containing short-chain fatty acids, sphingolipids containing very-long-chain fatty acids (≥20 carbons) exhibited the greatest increase in *Atg5* KO oligodendrocytes (Fig. 5J). In contrast, overall phospholipid levels, which follow different synthesis pathways from sphingolipids, remained largely unchanged in *Atg5* KO oligodendrocytes (fig. S6). The total levels of cholesterol, a major lipid component of myelin sheaths, remained unchanged in *Atg5* KO oligodendrocytes (Fig. 5K). To test whether ER-phagy restricts myelin sheath growth through degrading sphingolipid synthases, we leveraged the *Cers2* floxed mouse line to genetically perturb sphingolipid biosynthesis in oligodendrocytes (*52*). *Cers2* encodes a rate-limiting enzyme critical for the biosynthesis of very-long-chain ceramide (Fig. 5B), which forms the backbone of sphingolipids including galactosylceramide, sulfatide, and sphingomyelin (*51*). To disrupt sphingolipid biogenesis in oligodendrocytes under conditions of either normal or impaired ER-phagy flux, we generated *Cers2* single conditional knockout (*Cnp-Cre; Cers2^F/F^*) and *Atg5; Cers2* double conditional knockout (DKO) mice (*Cnp-Cre; Atg5^F/F^; Cers2^F/F^*). Western blot analysis confirmed efficient knockout of Atg5 and Cers2 and disrupted ER-phagy flux in *in vitro* differentiated *Atg5; Cers2* DKO oligodendrocytes (fig. S7, A and B).

Next, we examined whether genetic deletion of Cers2, which is aberrantly enriched in *Atg5* KO oligodendrocytes, could reverse the hypermyelination phenotype observed in *Atg5* cKO mice. As expected, *Atg5* cKO mice exhibited significantly increased myelin sheath thickness in the corpus callosum at P21, revealed by a significant reduction in *g*-ratios compared with controls (Fig. 6, A and B and fig. S7, C and D; quantified in Fig. 6E). Intriguingly, P21 *Cers2* cKO mice (*Cnp-Cre; Cers2^F/F^*) were not hypomyelinated compared to controls (Fig. 6C and fig. S7E; quantified in Fig. 6F), suggesting that Cers2 is not required for myelin formation at this stage. This is consistent with previous reports showing that Cers2 mutants exhibit thinner myelin sheaths beginning from 6 weeks of age (*53*), and that genetic deletion of Cgt, another rate-limiting enzyme for sphingolipid biogenesis, does not cause developmental myelination phenotypes (*54, 55*).

**Fig. 6.**
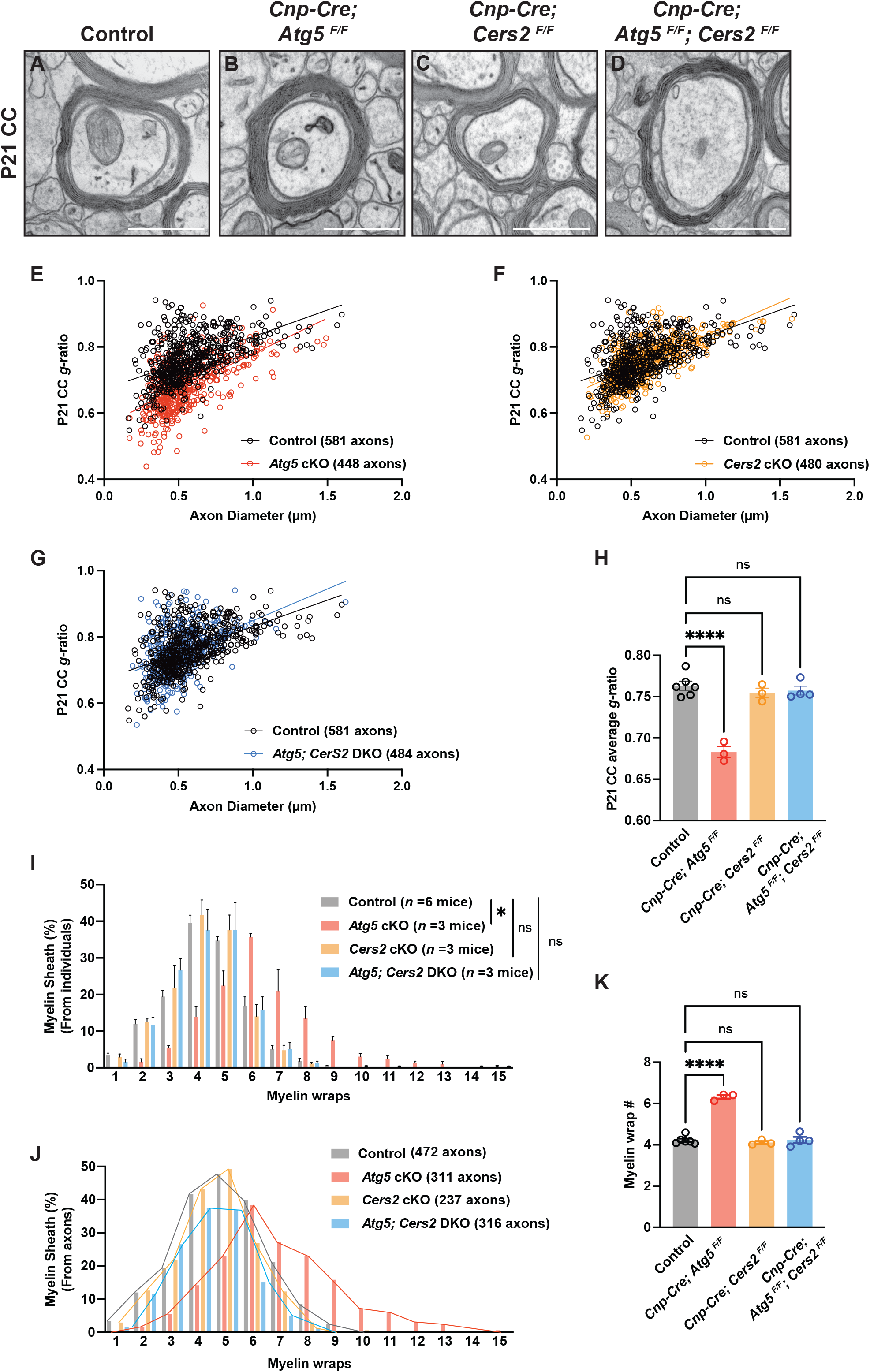
Genetic perturbation of sphingolipid biogenesis in the ER rescues hypermyelination observed in *Atg5* conditional knockout mice. **(A–D)** Representative TEM micrographs of corpus callosum (CC) from P21 control (*Atg5^F/F^* or *Atg5^F/F^; Cers2^F/F^*) (A), *Cnp-Cre; Atg5^F/F^* (B), *Cnp-Cre; Cers2^F/F^* (C), and *Cnp-Cre; Atg5^F/F^; Cers2^F/F^* mice (D). **(E–G)** Scatter plots showing *g*-ratios as a function of axon diameter in P21 *Cnp-Cre; Atg5^F/F^* (*Atg5* cKO) (E), *Cnp-Cre; Cers2^F/F^* (*Cers2* cKO) (F), and *Cnp-Cre; Atg5^F/F^; Cers2^F/F^* (*Atg5; Cers2* DKO) mice (G) compared with controls. **(H)** Quantification of average *g*-ratios in P21 control, *Cnp-Cre; Atg5^F/F^*, *Cnp-Cre; Cers2^F/F^*, and *Cnp-Cre; Atg5^F/F^; Cers2^F/F^*CC. **(I–K)** Quantification of the frequency distribution of myelin wrap numbers from individual mice (I), from all analyzed axons (J), and the average number of myelin wraps (K) among P21 control*, Atg5* cKO, *Cers2* cKO, and *Atg5; Cers2* DKO mice. One-way ANOVA followed by Tukey’s multiple comparisons test for (H) and (K). Kolmogorov-Smirnov test for (I). Scale bar: 0.5 µm in (A)–(D).

While *Cers2* cKO mice showed no change in myelin thickness at P21, *Atg5; Cers2* DKO mice reversed the hypermyelination phenotype observed in *Atg5* cKO mice (Fig. 6D and fig. S7F; quantified in Fig. 6, G and H). This is corroborated by the observation that both the distribution of myelin wrap numbers and average myelin wrap number per animal were rescued in *Atg5; Cers2* DKO mice compared to *Atg5* cKO (Fig. 6, I to K). Importantly, *Atg5; Cers2* DKO mice harbored similar numbers of oligodendrocytes in the cortex and cerebellum compared to *Atg5* cKO mice (fig. S8), showing that the rescue of myelin sheath thickness in *Atg5; Cers2* DKO was not due to changes in oligodendrocyte number. Thus, although Cers2 is not required for myelin formation during development, its excessive accumulation, due to the lack of ER-phagy, drives abnormal myelin thickening. Taken together, our results demonstrate that oligodendrocyte ER-phagy degrades sphingolipid synthases enriched in ER tubules, thereby reducing *de novo* sphingolipid biogenesis and suppressing myelin sheath growth (Fig. 7).

**Fig. 7.**
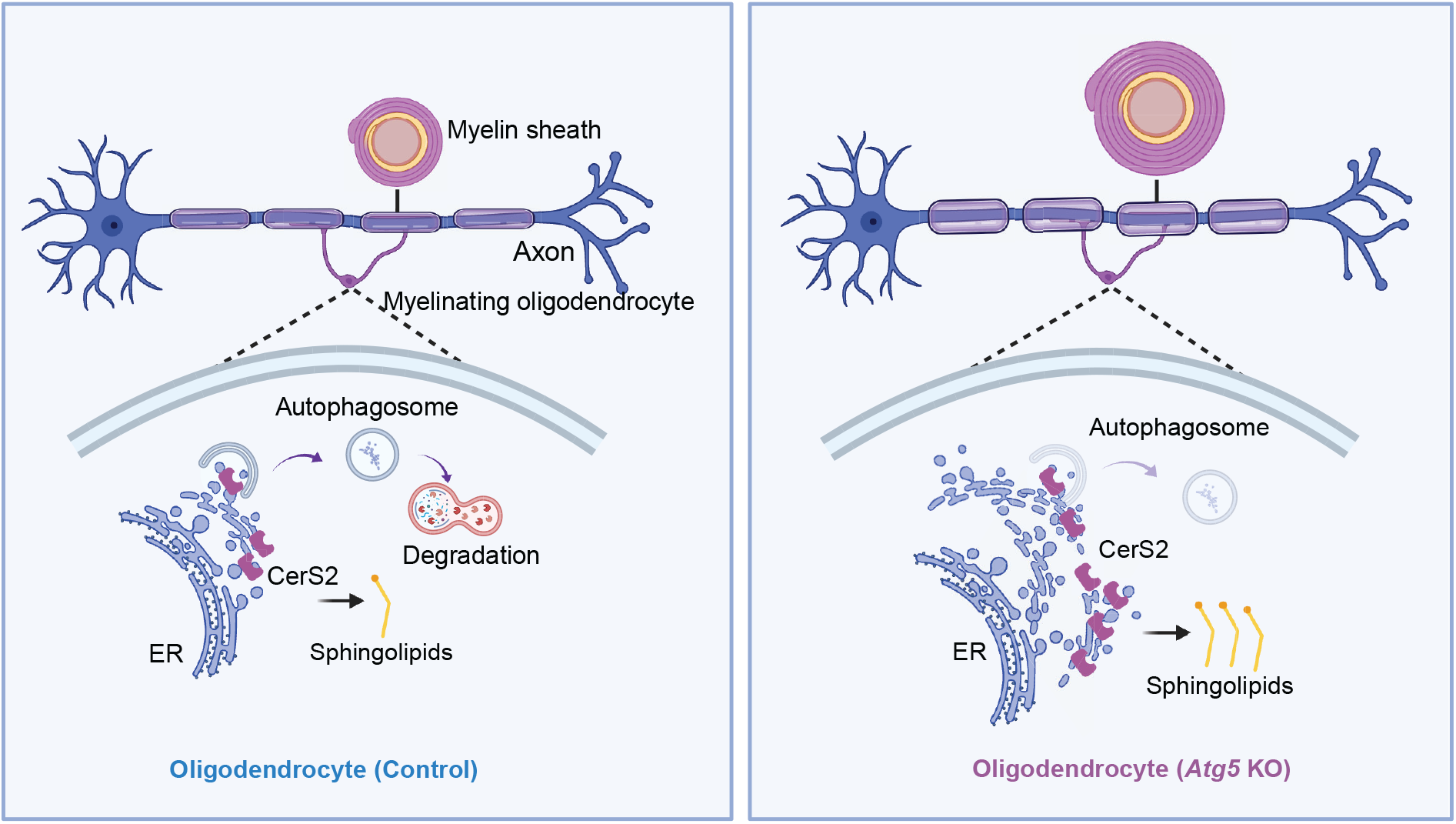
Oligodendrocyte ER-phagy degrades sphingolipid synthases located in ER tubules, thereby preventing excessive myelin sheath growth. In WT oligodendrocytes, autophagy degrades portions of the ER (ER-phagy) to limit the abundance of ER-tubule–resident sphingolipid biosynthetic enzymes (e.g., Cers2), thereby restricting sphingolipid levels and ensuring proper myelin sheath thickness. Perturbation of autophagy in oligodendrocytes leads to ER accumulation, increased levels of sphingolipid synthases, and elevated sphingolipid production. This further results in excessive myelin sheath growth and increased sheath thickness.

## DISCUSSION

Myelin formation during CNS development is precisely regulated to ensure proper axonal insulation and neural circuit function. Previous studies have shown that IGF receptor activation, sustained Akt signaling, *Pten* deletion, *Tfeb* inactivation, ErbB pathway activation, or persistent Erk1/2 activation can all lead to increased myelin sheath thickness (*7–13, 16, 22, 23*). Here, we reveal a distinct mechanism that restricts myelin sheath growth: ER-phagy acts as an oligodendrocyte-intrinsic “brake” on myelin formation. This is achieved through the selective degradation of ER tubules and resident lipid synthases, thereby limiting sphingolipid production and repressing myelin sheath growth.

Previous studies have largely positioned autophagy in oligodendrocytes as a homeostatic mechanism operating in the mature or aging CNS, where it preserves myelin integrity through turnover of myelin proteins (*31, 34, 36*). Our previous work has shown that autophagy collaborates with apoptosis pathways to limit the survival of premyelinating oligodendrocytes during early brain development, demonstrating its critical role in oligodendrocyte lineage progression (*30*). Our present study reveals a fundamentally distinct and temporally specific role of autophagy: during developmental myelination, oligodendrocyte autophagy powerfully represses myelin membrane growth, thereby actively constraining sheath thickness rather than merely sustaining cell viability or maintaining existing myelin. Mechanistically, ER-phagy–dependent control of sphingolipid biosynthesis serves as the link between selective autophagy and myelin membrane biogenesis, revealing the molecular underpinning of this “brake” on myelin sheath growth.

ER-phagy, a selective form of autophagy that targets the ER for degradation, has recently emerged as an essential process regulating nervous system homeostasis (*56*). For instance, degradation of ER tubules has been observed in mouse brain axons and human induced pluripotent stem cell (iPSC)-derived neurons (*27–29*), where it plays a critical role in presynaptic neurotransmission by regulating calcium release from ER stores via ryanodine receptors (*28*). A recent study reported that ER tubules extend into the inner tongue of myelin sheaths, mediating nonvesicular lipid transfer required for myelination (*57*). Consistent with the positive role of ER tubules in promoting myelin growth, our study demonstrates that ER-phagy decreases ER tubules and sphingolipid biogenesis in oligodendrocytes, thereby restricting myelin production. It will be of great interest to investigate the extent to which ER-phagy occurs in the inner tongue of the myelin sheath to locally regulate myelin sheath remodeling.

Several mammalian ER-phagy receptors have been identified to date, including Fam134a/b/c, Atl3, Ccpg1, Tex264, Calcoco1, and Rtn3 (*26*). Notably, redundancy among these receptors has been reported in heterologous cells and neurons, as knockout of individual ER-phagy receptors often produces minimal phenotypic effects (*27, 28*). In our study, we detected significant enrichment of multiple ER-phagy receptors in *Atg5* knockout oligodendrocytes (Fig. S3), suggesting functional redundancy among these receptors in oligodendrocytes. Moreover, it remains unclear whether ER degradation in oligodendrocytes is mediated by known ER-phagy receptors or by yet-unidentified mechanisms. Together, these observations present a major challenge for identifying the ER-phagy receptor(s) responsible for ER tubule turnover in oligodendrocytes, and future studies employing advanced genetic and imaging approaches will be necessary to address this question.

Sphingolipids are major structural components of myelin, accounting for approximately 25% of total myelin lipids (*24*). Oligodendrocytes rely heavily on intrinsic sphingolipid biosynthesis, particularly sphingolipids containing very-long-chain fatty acids (*24, 34*). Previous studies have demonstrated that oligodendrocyte-specific deletion of Cers2, an enzyme responsible for synthesizing very-long-chain sphingolipids, leads to myelin degeneration in adult mice (*53*). Moreover, global *Cers2* knockout causes myelin abnormalities in trigeminal nerve axons (*58*). These findings highlight the critical role of very-long-chain sphingolipids in maintaining myelin integrity. Our study shows that *Cers2* cKO mice exhibit no obvious changes in myelin integrity or myelin wrap numbers at P21 (Fig. 6), but excessive accumulation of Cers2 due to ER-phagy perturbation causes hypermyelination (Fig. 1, 2, and 6). This is consistent with previous studies showing that Cers2 deletion does not result in an absence of sphingolipids, but rather in a shift from very-long-chain sphingolipids to short-chain sphingolipids (*53, 58*). Additionally, our results showing that Cers2 is not required for the establishment of myelination are corroborated by prior studies demonstrating that Cgt, another rate-limiting enzyme in sphingolipid biogenesis, is dispensable for developmental myelination but essential for myelin maintenance (*54, 55*). Therefore, the relatively normal myelination observed in juvenile *Cers2* mutants suggests that Cers2-deficient oligodendrocytes may be resilient to the loss of very-long-chain sphingolipids during early brain development, possibly through compensatory mechanisms activated upon Cers2 deletion.

Our data show that *Atg5* cKO oligodendrocytes exhibit elevated sphingolipids, and that deletion of *Cers2* in the *Atg5* mutant background restores myelin wrap numbers and sheath thickness to normal levels observed in *Atg5; Cers2* DKO mice (Fig. 6). This suggests that the accumulation of very-long-chain sphingolipids in *Atg5*-deficient oligodendrocytes is sufficient to promote myelin sheath growth and wrapping. However, how do elevated sphingolipids drive hypermyelination? Previous studies have shown that, apart from being essential myelin lipids, sphingolipids such as galactosylceramide are critical for the proper assembly and trafficking of myelin proteins such as myelin proteolipid protein (Plp) (*59*). Therefore, elevated very-long-chain sphingolipids in Atg5-deficient oligodendrocytes may enhance Plp delivery and incorporation into the myelin membrane, thereby increasing myelin sheath formation observed in *Atg5* cKO mice. Future studies are needed to test this hypothesis.

In conclusion, our findings establish ER-phagy as a cellular brake on developmental myelination and uncover that the ER-phagy-sphingolipid axis restricts myelin membrane expansion. Together, these results reveal a mechanistic link between ER homeostasis and myelin production, providing a foundation for understanding the mechanisms and pathological consequences of hypermyelination.

## ACKNOWLEDGMENTS

We thank Drs. Klaus-Armin Nave and Jens Brüning for sharing the *Cnp-Cre* and *Cers2* floxed mice, respectively. We thank all the members of the Sun laboratory for assistance and discussions. We thank the UT Southwestern Electron Microscopy Core Facility, especially Dr. Natalia Gunko, Phoebe Doss, Ethan Ozment, and Jaryd Gipson, for their help with TEM sample processing; the UT Southwestern Proteomics Core Facility, especially Dr. Andrew Lemoff, for proteomic analyses; the UT Southwestern Quantitative Light Microscopy Core Facility, especially Dr. Marcel Mettlen, for assistance with confocal imaging; and the UT Medical Branch Mass Spectrometry Facility, especially Dr. William Russell, Trevor Romsdahl, and Jennifer Linares, for lipidomic analyses.

## Author contributions

Conceptualization, methodology, investigation, writing, and funding acquisition: T.Z. and L.O.S.; Methodology and investigation: D.B.; and supervision: L.O.S.

## SUPPLEMENTAL MATERIALS

### Supplemental Figures

**Fig. S1, related to Fig. 1.**
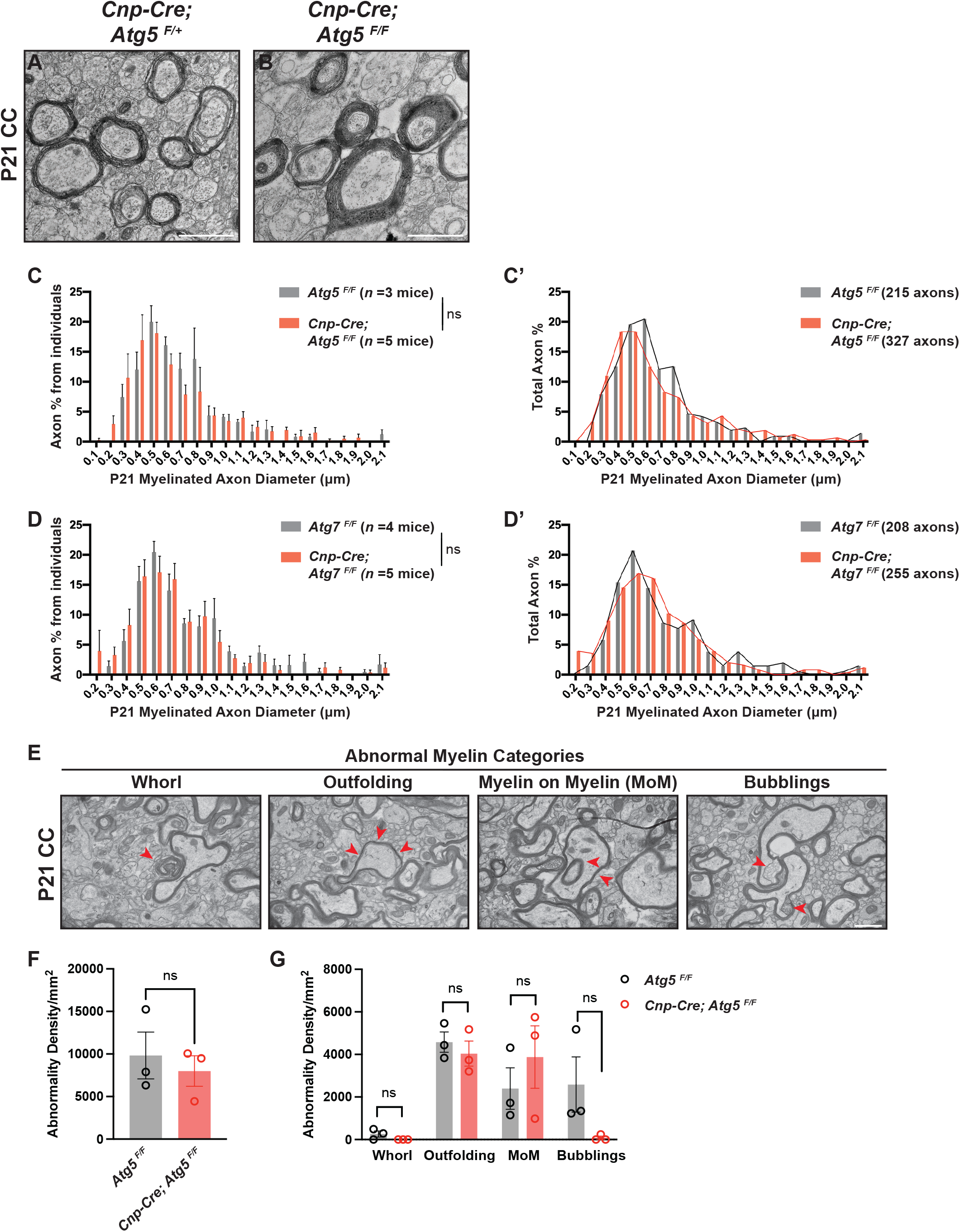
Characterization of axon diameter and myelin abnormalities in autophagy-deficient oligodendrocyte conditional knockout mice. **(A and B)** Representative TEM micrographs from P21 *Cnp-Cre; Atg5^F/+^* (A) and *Cnp-Cre; Atg5^F/F^* (B) corpus callosum (CC). **(C–D’)** Frequency distribution of axons with different diameters in P21 *Atg5^F/F^* and *Cnp-Cre; Atg5^F/F^* corpus callosum (C–C’), and in P21 *Atg7^F/F^*and *Cnp-Cre; Atg7^F/F^* corpus callosum (D–D’). **(E)** Representative TEM micrographs from P21 *Atg5^F/F^* mice showing different myelin abnormalities. The abnormal myelin structures were indicated by red arrowheads. **(F and G)** Quantification of the density of total abnormal myelin profiles (F) and individual myelin abnormalities (G) in P21 *Atg5^F/F^* and *Cnp-Cre; Atg5^F/F^* corpus callosum. Kolmogorov–Smirnov test for (C) and (D). Two-tailed *t*-test for (F) and (G). Scale bars: 1 μm in (A), (B), and (E).

**Fig. S2, related to Fig. 2.**
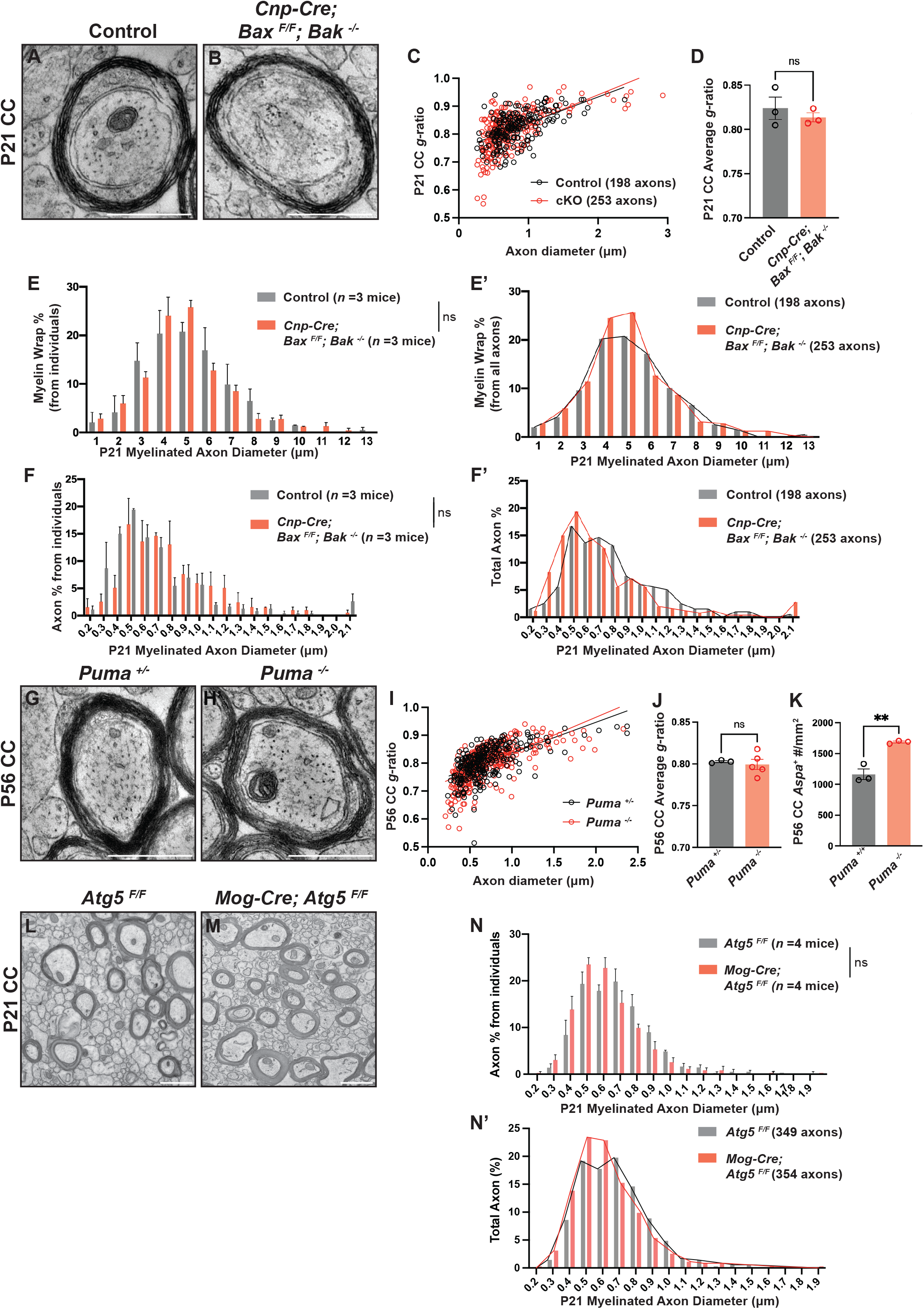
Characterization of myelin sheath thickness and axon diameter distributions in *Cnp-Cre; Bax^F/F^; Bak^-/-^*, *Puma^-/-^*, and *Mog-Cre; Atg5^F/F^* mice. **(A and B)** Representative TEM micrographs of P21 corpus callosum from *Cnp-Cre; Bax^F/F^; Bak^-/-^* mutants (B) and littermate controls (*Cnp-Cre; Bax^F/+^; Bak^+/-^*, A). **(C)** Scatter plot showing *g*-ratios of myelinated axons as a function of axon diameter in P21 *Cnp-Cre; Bax^F/F^; Bak^-/-^* mutants (red) and littermate controls (*Cnp-Cre; Bax^F/+^; Bak^+/-^*, black). **(D)** Quantification of average *g*-ratios of myelinated axons in P21 *Cnp-Cre; Bax^F/F^; Bak^-/-^* mutants and littermate controls (*Cnp-Cre; Bax^F/+^; Bak^+/-^*). **(E–F’)** Frequency distribution of axons with different myelin wrap numbers (E and E’), and frequency distribution of axons with different diameters (F and F’) in the corpus callosum from P21 *Cnp-Cre; Bax^F/F^; Bak^-/-^* mutants and littermate controls. **(G and H)** Representative TEM micrographs of P56 *Puma^+/-^* (G) and *Puma^-/-^* (H) corpus callosum axons. **(I and J)** Quantification of average *g*-ratios of myelinated axons (J), and as a function of axon diameter (I), in P56 *Puma^-/-^*mutants (red) compared to littermate controls (black). *n* ≥ 200 axons from at least 3 animals per genotype in I, and *n* ≥ 3 animals for each genotype in J. **(K)** Quantification of *Aspa^+^* mature oligodendrocyte numbers in the corpus callosum of P56 *Puma^+/+^* and *Puma^-/-^* mice. **(L and M)** Representative TEM micrographs from P21 *Atg5^F/F^* (L) and *Mog-Cre; Atg5^F/F^* (M) corpus callosum. **(N and N’)** Frequency distribution of axons with different diameters in P21 *Atg5^F/F^* and *Mog-Cre; Atg5^F/F^* corpus callosum. Kolmogorov–Smirnov test for (E), (F), and (N). Two-tailed *t*-test for (D), (J), and (K). Scale bars: 0.5 μm in (A), (B), (G), and (H); 1 μm in (L) and (M).

**Fig. S3, related to Fig. 3.**
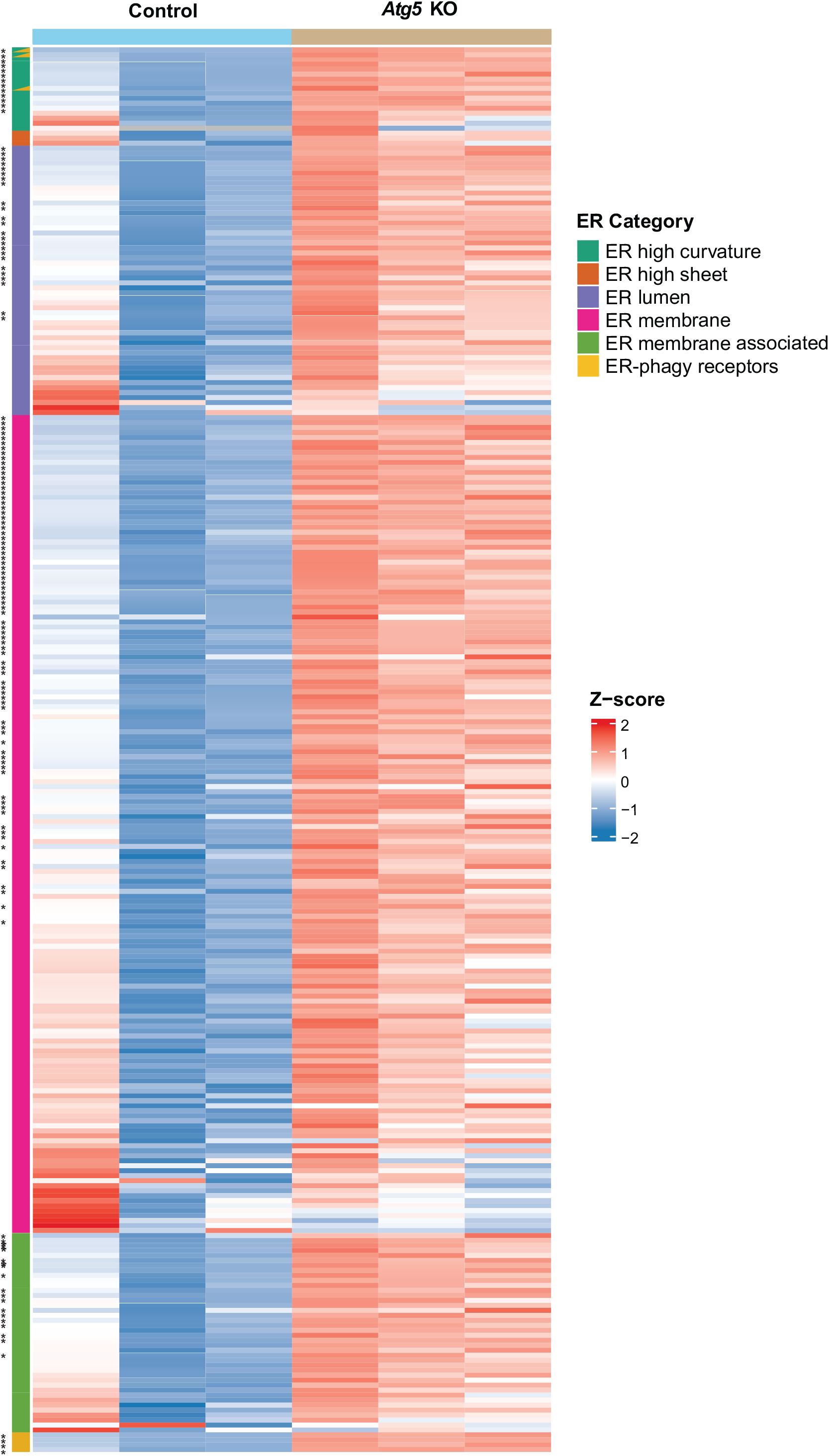
Characterization of ER-related protein expression in *Cnp-Cre; Atg5^F/F^* oligodendrocytes. Heatmap showing relative protein levels (z-scores) of ER proteins belonging to different categories in control (*Atg5^F/F^*) and *Atg5* KO *(Cnp-Cre; Atg5^F/F^)* oligodendrocytes, as determined by quantitative proteomics. Each column represents cells purified from an individual mouse. Each row represents a protein. Proteins assigned to two ER-related categories are shown as diagonally split, two-colored rectangles in the left annotation bar. Significantly increased proteins (*p* < 0.05) are indicated by asterisks.

**Fig. S4, related to Fig. 3.**
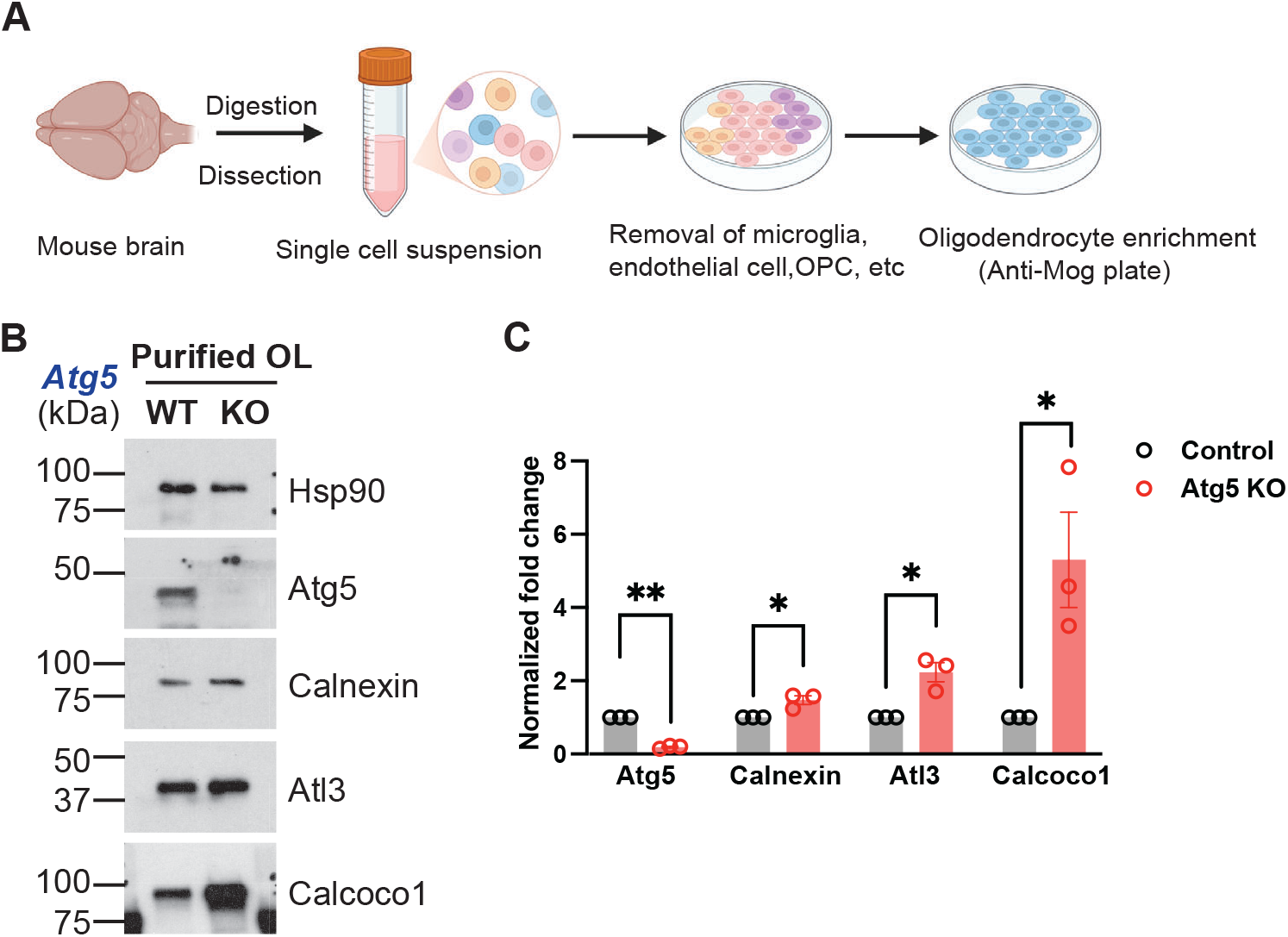
Acutely purified oligodendrocytes from *Cnp-Cre; Atg5^F/F^* mice exhibit a significant elevation in ER-resident protein levels. **(A)** Workflow of purifying mature oligodendrocytes from mouse brains (created with BioRender). **(B)** Western blots showing the protein levels of Atg5, Calnexin, and ER-phagy receptors Atl3 and Calcoco1 in purified oligodendrocytes from *Cnp-Cre; Atg5^F/F^* and *Atg5^F/F^* mice. **(C)** Normalized fold change of indicated protein levels in purified oligodendrocytes from *Cnp-Cre; Atg5^F/F^* and *Atg5^F/F^* mice. Protein levels were normalized to Hsp90. *n* = 3 independent cell purifications. One-sample *t*-test for (C).

**Fig. S5, related to Fig. 4.**
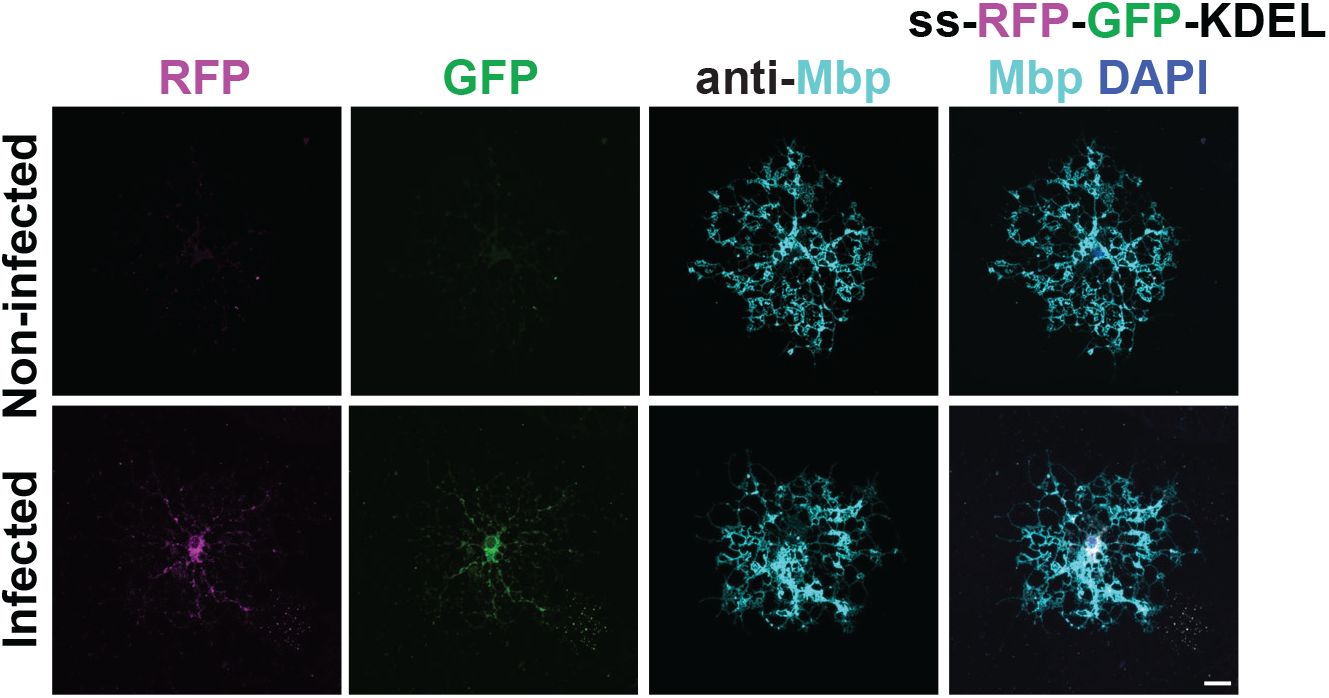
Characterization of lentiviral expression of an ER-phagy reporter in oligodendrocytes. Representative confocal images of oligodendrocytes that were either uninfected (top) or transduced with the ER-phagy reporter (ss-RFP-GFP-KDEL; bottom). The cells were immunostained for Mbp (cyan), with simultaneous detection of native RFP and GFP fluorescence. Scale bar: 20 μm.

**Fig. S6, related to Fig. 5.**
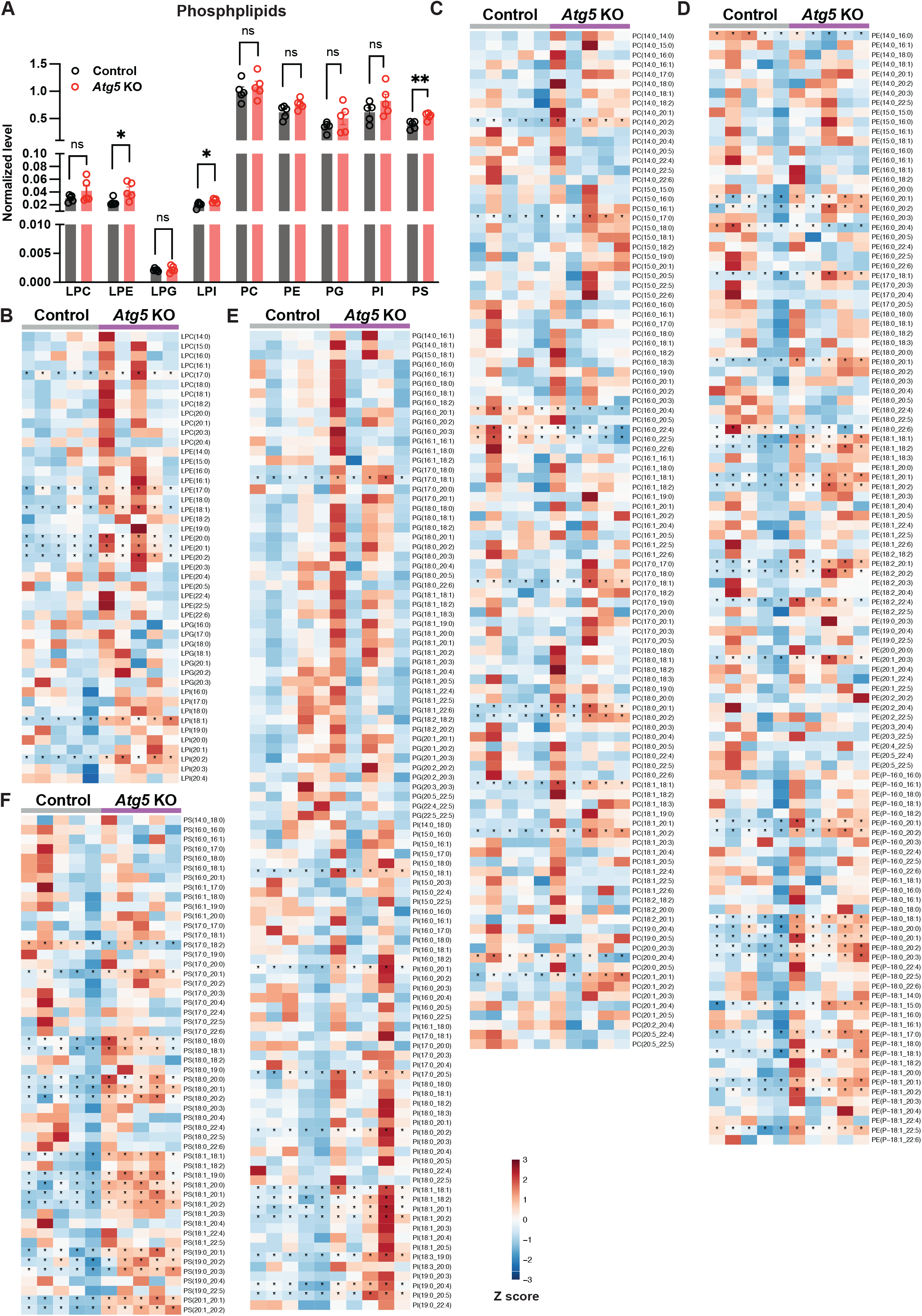
Characterization of phospholipid levels in *Cnp-Cre; Atg5^F/F^* oligodendrocytes. **(A)** Quantification of normalized phospholipid levels in *Atg5* KO (*Cnp-Cre; Atg5^F/F^*) compared with control (*Atg5^F/F^*) oligodendrocytes using lipidomics. Normalization was based on lipid reads per total protein level. *n* = 5 independent cell purifications. LPC, lysophosphatidylcholine; LPE, lysophosphatidylethanolamine; LPG, lysophosphatidylglycerol; LPI, lysophosphatidylinositol; PC, phosphatidylcholine; PE, phosphatidylethanolamine; PG, phosphatidylglycerol; PI, phosphatidylinositol; PS, phosphatidylserine. Two-tailed *t*-test for (A). **(B–F)** Heatmaps showing relative levels (z-scores) of phospholipid species with varying fatty acid chains from control (*Atg5^F/F^*) and *Atg5* KO (*Cnp-Cre; Atg5^F/F^*) oligodendrocytes at day 8 *in vitro*, revealed by lipidomics. Significantly increased lipids (*p* < 0.05) are indicated by asterisks. LPC, LPE, LPG, and LPI are shown in (B); PC in (C); PE in (D); PG and PI in (E); and PS in (F).

**Fig. S7, related to Fig. 6.**
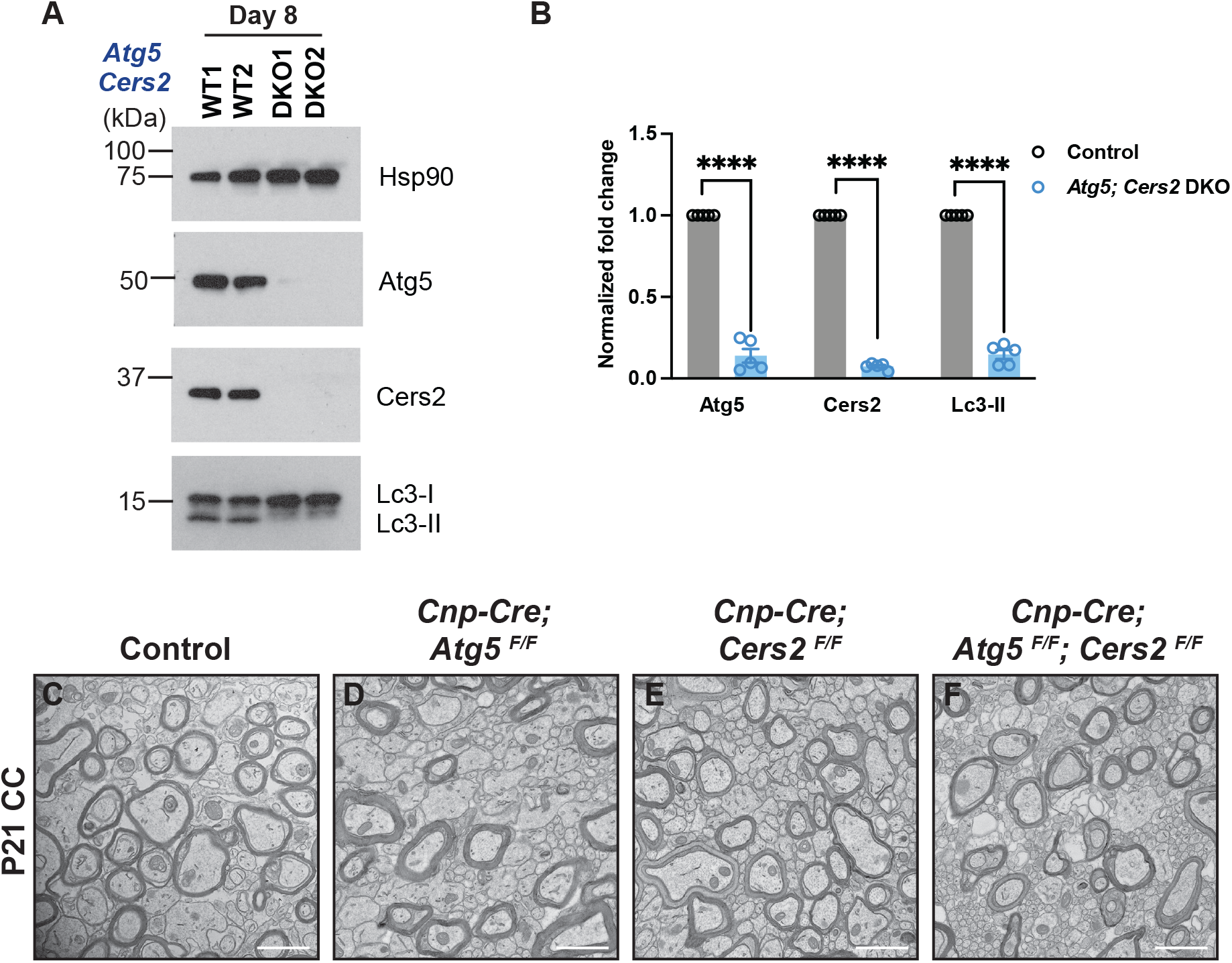
Characterization of protein expression, autophagy flux, and myelination in *Cnp-Cre; Atg5^F/F^; Cers2^F/F^* double conditional knockout mice. **(A)** Western blots showing Atg5, Cers2, and Lc3 protein levels in WT (*Atg5^F/F^; Cers2^F/F^*) and *Atg5; Cers2* DKO *(Cnp-Cre; Atg5^F/F^; Cers2^F/F^*) oligodendrocytes at day 8 *in vitro*. **(B)** Normalized fold change of indicated protein levels in *Atg5; Cers2* DKO oligodendrocytes compared with WT oligodendrocytes at day 8 *in vitro*. Protein levels were normalized to Hsp90. *n* = 5 independent cell purifications. **(C*–*F)** Representative TEM micrographs of corpus callosum (CC) from P21 control (*Atg5^F/F^* or *Atg5^F/F^; Cers2^F/F^*) (C), *Cnp-Cre; Atg5^F/F^* (D), *Cnp-Cre; Cers2^F/F^* (E), and *Cnp-Cre; Atg5^F/F^; Cers2^F/F^* mice (F). One-sample *t*-test for (B). Scale bars: 1 μm in (C)***–***(F).

**Fig. S8, related to Fig. 6.**
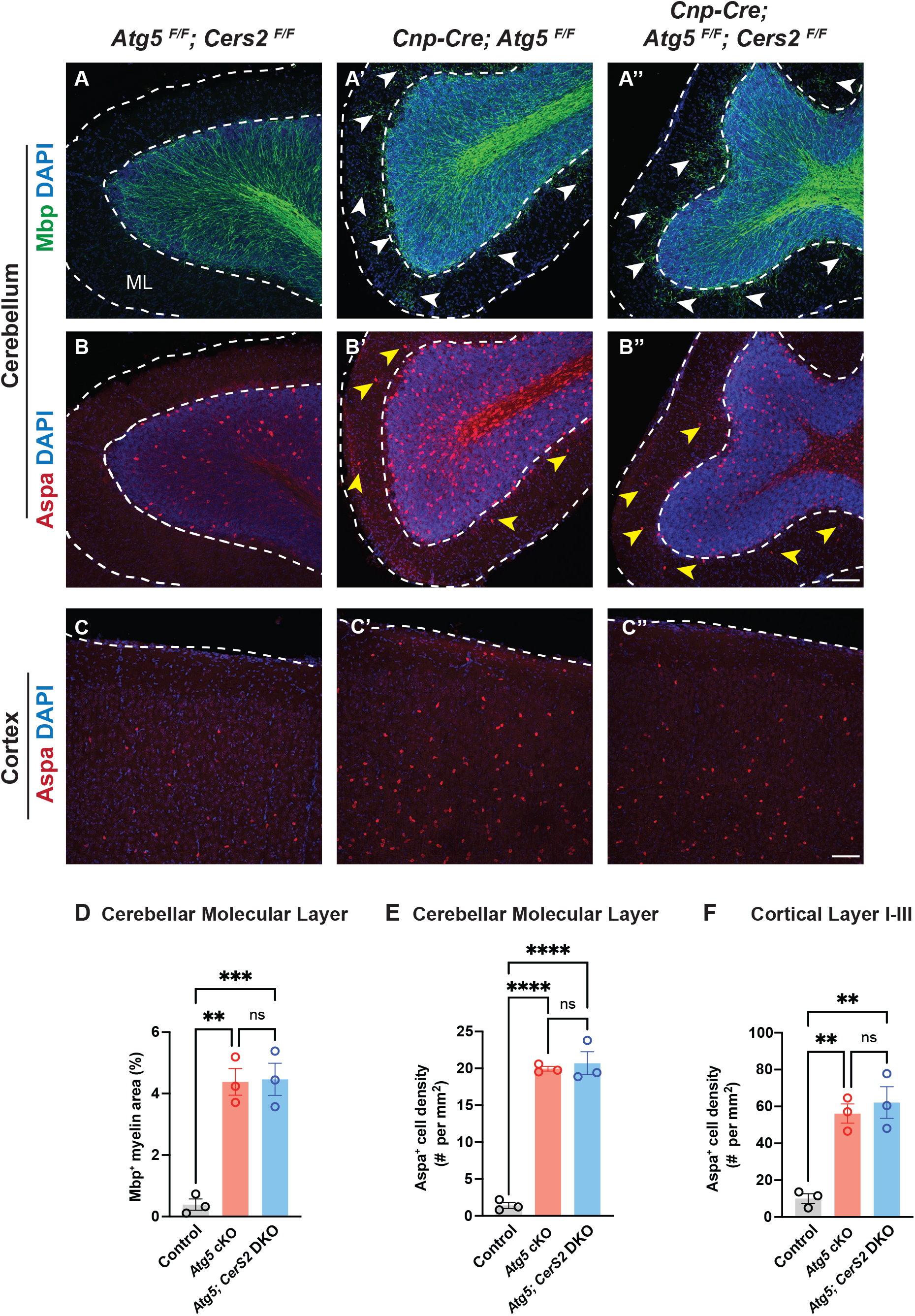
Oligodendrocyte numbers and ectopic myelination of the cerebellar molecular layer in *Cnp-Cre; Atg5^F/F^; Cers2^F/F^* mice are comparable to those in *Cnp-Cre; Atg5^F/F^* mice. **(A–B”)** Sagittal brain sections from P21 mice showing ectopic myelination in the cerebellar molecular layer (ML), indicated by aberrant Mbp fluorescence (white arrowheads in A’ and A”) and Aspa^+^ mature oligodendrocytes (yellow arrowheads in B’ and B”) in *Cnp-Cre; Atg5^F/F^* mice (A’ and B’) and *Cnp-Cre; Atg5^F/F^; Cers2^F/F^ mice* (A” and B”), as compared with *Atg5^F/F^; Cers2^F/F^* mice (A and B). Dashed lines demarcate the cerebellar molecular layer. **(C–C”)** Sagittal brain sections showing Aspa^+^ oligodendrocytes in the cortex of *Atg5^F/F^; Cers2^F/F^* (C), *Cnp-Cre; Atg5^F/F^* (C’), and *Cnp-Cre; Atg5^F/F^; Cers2^F/F^* mice (C”). Dashed lines indicate the pia surface of the cortex. **(D–F)** Quantification of Mbp^+^ myelin area percentage in the cerebellar molecular layer (D), Aspa^+^ oligodendrocyte numbers in the cerebellar molecular layer (E), and Aspa^+^ oligodendrocyte numbers in cortical layer I–III (F) of *Atg5^F/F^; Cers2^F/F^*, *Cnp-Cre; Atg5^F/F^, and Cnp-Cre; Atg5^F/F^; Cers2^F/F^* mice. One-way ANOVA followed by Tukey’s multiple comparisons test for (D)–(F). Scale bars: 100 µm in (B”) for (A)–(B”); and 100 µm in (C”) for (C)–(C”).

### Other Supplemental Items

**Table S1: Quantitative proteomic analysis of dysregulated proteins in *in vitro* differentiated *Cnp-Cre; Atg5^F/F^* (*Atg5* KO) oligodendrocytes as compared to *Atg5^F/F^*(WT) oligodendrocytes, related to Fig. 3**.

**Table S2: Lipidomic analysis of sphingolipids and phospholipids in *in vitro* differentiated *Cnp-Cre; Atg5^F/F^* (*Atg5* KO) oligodendrocytes as compared to *Atg5^F/F^*(WT) oligodendrocytes, related to Fig. 5**.

## MATERIALS AND METHODS

### Animals

All animal procedures were carried out in accordance with NIH guidelines and were approved by the Institutional Animal Care and Use Committee (IACUC) at the University of Texas Southwestern Medical Center. Mice were maintained under standard 12-hour light/12-hour dark cycle with food and water provided *ad libitum*. The day of birth was defined as postnatal day 0 (P0), and developmental stages are indicated in the figures and legends. Both male and female mice were included in all experiments and analyses. Transgenic mouse lines used in this study included *Atg5^Flox^* (*Atg5^F^*, RIKEN RBRC02975), *Atg7^Flox^* (*Atg7^F^*, RIKEN RBRC02759), *Cnp-Cre* (*38*), *Mog-Cre* (JAX#041041) (*22*), *Cers2^Flox^* (*Cers2^F^*) (*52*), *Puma^-^* (JAX#011067) (*23*), and *Bax^Flox^; Bak^-^* (*Bax^F^; Bak^-^*, JAX#006329).

### Isolation of mouse oligodendrocyte precursor cells (OPCs), *in vitro* differentiation, and pharmacological treatments

OPCs were purified from postnatal day 6–10 mouse brains using an immunopanning-based approach adapted from previously published protocols (*30, 60*). Briefly, pups were rapidly decapitated, and whole brains (excluding olfactory bulbs) were dissected and minced into ∼1 mm³ pieces. Tissue fragments were enzymatically dissociated in a digestion buffer consisting of 1× Earle’s Balanced Salt Solution (10 × EBSS; Sigma-Aldrich, Cat#E7501), 1 mM MgSO₄, 0.46% glucose, 2 mM EGTA, 26 mM NaHCO₃, 20 U/mL papain (Worthington, Cat#LS003126), and 250 U/mL DNase I (Worthington, Cat#LS002007). Digestion was carried out at 34°C for 90 minutes under 5% CO₂/95% O₂ perfusion. Following enzymatic treatment, tissues were triturated with 1 mL pipettes to obtain a single-cell suspension. The suspension was sequentially incubated on three BSL-1–coated Petri dishes (Vector Laboratories, Cat#L1100) for 10 minutes each to deplete endothelial cells and microglia. OPCs were then selectively captured on plates coated with rat anti-Pdgfrα antibody (BD Biosciences, Cat#558774).

Purified OPCs were cultured in proliferation medium composed of DMEM-SATO base medium supplemented with 4.2 μg/mL forskolin (Sigma-Aldrich, Cat#F6886), 10 ng/mL PDGF-AA (Peprotech, Cat#100-13A), 10 ng/mL CNTF (Peprotech, Cat#450-02), and 1 ng/mL NT-3 (Peprotech, Cat#450-03) (*60*). Cells were maintained at 37°C in a humidified incubator with 10% CO₂. Differentiation was induced by replacing the proliferation medium with differentiation medium (DMEM-SATO supplemented with 4.2 μg/mL forskolin, 10 ng/mL CNTF, and 40 ng/mL T3; Sigma-Aldrich, Cat#T6397). Half of the culture medium was refreshed every 48 hours. The oligodendrocytes at differentiation day 8 were used for experiments.

For pharmacological manipulations, oligodendrocytes were treated with either 5 nM bafilomycin A1 (BafA1; Sigma-Aldrich, Cat#196000), 1 μM VPS34IN1 (Selleckchem, Cat#S7980), or an equivalent volume of DMSO (vehicle control) for 24 hours at 37°C.

### Isolation of mouse oligodendrocytes

Oligodendrocytes were purified from postnatal day 13–14 mouse brains using an immunopanning method adapted from previously described protocols (*60*). Tissue digestion and preparation of single-cell suspensions were performed as described for mouse OPC isolation. The cell suspension was sequentially incubated on two BSL-1–coated Petri dishes for 10 minutes each, followed by incubation on a rat anti-CD45 antibody–coated dish (BD Biosciences, Cat#550539) for 20 minutes to deplete endothelial cells and microglia. The suspension was then transferred to dishes coated with rat anti-Pdgfrα antibody (45 minutes) and anti-A2B5 antibody ascites (30 minutes) to deplete OPCs. Mature oligodendrocytes were subsequently enriched by positive selection on a dish coated with anti-Mog hybridoma supernatant.

### Immunostaining

For mouse tissue immunostaining, mice were euthanized with CO₂ and perfused with ice-cold PBS followed by 4% PFA in PBS. Brains were post-fixed in 4% PFA at 4°C overnight, cryoprotected in 30% sucrose for 24 hours, and sectioned at 40 µm thickness with a microtome. Brain sections were pretreated with L.A.B (Liberate Antibody Binding) solution (Polysciences, Cat#24310) for 10 minutes and then blocked in PBST (PBS containing 0.5% Triton X-100) with 10% donkey serum for 20 minutes and incubated overnight at 4°C with primary antibodies in PBST containing 2% donkey serum. After three washes in PBST, sections were incubated for 1 hour at room temperature with Alexa Fluor 488- or 555-conjugated secondary antibodies (1:1000; Thermo Fisher Scientific) and DAPI (0.1 µg/mL; Thermo Fisher Scientific). Sections were washed and mounted on glass slides using VectaShield Hardset Antifade medium (Vector Laboratories, Cat#H- 1400-10). Fluorescent images were acquired using a Keyence BZ-810 all-in-one fluorescence microscope or a Nikon AXR inverted confocal microscope. For quantification of Mbp-positive areas and Aspa-positive cells, entire brain sections were captured using the automated image-stitching function of the Keyence BZ-810. Stitched images were analyzed using ImageJ software. Primary antibodies used in this study include rat anti-Mbp (1:4000, Abcam, Cat#ab7349) and rabbit anti-Aspa (1:1000, GeneTex, Cat#GTX113389; or 1:200, Sigma-Aldrich, Cat# ABN1698).

For immunostaining of cultured oligodendrocytes, OPCs were cultured and differentiated for 8 days on poly-D-lysine–coated coverslips. The cells were washed with PBS and fixed with 4% PFA for 20 minutes at room temperature. Following fixation, cells were incubated in 100 mM glycine in PBS for 10 minutes to quench residual aldehydes and reduce background fluorescence. Cells were then permeabilized and blocked with 10% donkey serum in PBST (PBS containing 0.1% Triton X-100 in) for 20 minutes, followed by an additional blocking step in PBS containing 2.5% BSA for 20 minutes. Primary antibody incubation was performed overnight at 4°C with rat anti-Mbp (1:500, Bio-Rad, Cat#MCA409S) diluted in PBS containing 2% donkey serum and 1% BSA. After three washes with PBS, cells were incubated for 1 hour at room temperature with Alexa Fluor 647–conjugated donkey anti-rat IgG secondary antibody (1:1000; Thermo Fisher Scientific, Cat#A78947) and DAPI (0.1 µg/mL; Thermo Fisher Scientific). Cells were washed and mounted onto glass slides using ProLong Diamond Antifade Mountant (Thermo Fisher Scientific, Cat#P36970). Fluorescent images were acquired using a Nikon AXR inverted confocal microscope.

### Western immunoblotting

For oligodendrocyte *in vitro* differentiation, OPCs were cultured either in 6-well plates (100,000 seeding cells per well) or in 6-cm dishes (250,000 seeding cells per dish). For acutely purified oligodendrocytes, cells were enriched by positive selection on anti-Mog hybridoma supernatant–coated Petri dishes and subsequently dissociated with Accutase cell detachment solution (STEMCELL Technologies, Catalog#07922). Cells were rinsed with ice-cold PBS and lysed in RIPA buffer (Thermo Fisher Scientific, Cat#89900) supplemented with 1× cOmplete protease inhibitor cocktail (Sigma-Aldrich, Cat#5892791001) and 1× PhosSTOP phosphatase inhibitor cocktail (Sigma-Aldrich, Cat#4906845001). Lysates were incubated for 1 hour at 4°C and centrifuged at 14,000 rpm for 10 minutes. Supernatants were harvested for further analysis.

Protein concentrations were determined using the BCA Protein Assay Kit (Thermo Fisher Scientific, Cat#23227). Samples were denatured at 95°C for 10 minutes with 4× LDS sample buffer (Thermo Fisher Scientific, Cat#NP0007) containing 10% 2-mercaptoethanol (Sigma-Aldrich, Cat# M6250). 1–5 µg protein was loaded onto 4–20% stain-free gradient precast polyacrylamide gels (Bio-Rad, Cat#4568096) and separated by SDS-PAGE. Proteins were then transferred to 0.45 µm or 0.22 µm PVDF membranes, and the membranes were blocked with EveryBlot Blocking Buffer (Bio-Rad, Cat#12010020) for at least 20 minutes at room temperature. The membranes were then incubated overnight at 4°C on a rocker with primary antibodies diluted in EveryBlot Blocking Buffer. After washing three times in TBST (TBS buffer with 0.1% Tween-20, 10 minutes each), membranes were incubated with HRP-conjugated secondary antibodies diluted in 5% non-fat milk for 1 hour at room temperature. Blots were washed in TBST and developed using either Amersham ECL Prime (Cytiva, Cat#RPN2232) or SuperSignal West Femto Maximum Sensitivity Substrate (Thermo Fisher Scientific, Cat#34095) for 3 minutes. Signals were visualized on X-ray films. For quantification, films were scanned and analyzed using ImageJ software. Protein signals were normalized to Gapdh, β-actin, or Hsp90.

Primary antibodies used were: rabbit anti-Atg5 (1:1000, Cell Signaling Technology, Cat#12994), rabbit anti-p62 (1:1000, Cell Signaling Technology, Cat#5114), mouse anti-Lc3 (1:500, MBL, Cat#M186-3), mouse anti-Gapdh (1:500, Santa Cruz Biotechnology, Cat#sc-32233), mouse anti-β-actin (1:500, Santa Cruz Biotechnology, Cat#sc-47778), rabbit anti-Hsp90 (1:5000, Cell Signaling Technology, Cat#4877), rabbit anti-Ckap4 (1:1000, Proteintech, Cat#16686-1-AP), rabbit anti-Calnexin (1:10000, Abcam, Cat#ab213243), rabbit anti-Rtn3 (1:5000, Sigma-Aldrich, Cat# ABN1723), mouse anti-Sptlc1 (1:1000, Santa Cruz Biotechnology, Cat#sc-374143), and rabbit anti-Cers2 (1:2000, Sigma-Aldrich, Cat#HPA027262).

### Transmission electron microscopy imaging

Mice were perfused with PBS followed by a fixative solution containing 4% paraformaldehyde, 2.5% glutaraldehyde, and 0.2% picric acid in phosphate buffer. Brains were post-fixed in the same solution at 4°C for 24 hours before further dissection. For transmission electron microscopy (TEM) analysis of the corpus callosum (CC), coronal brain sections at 1 mm thickness near the midline were prepared, and only the medial region of the CC was dissected for processing. Samples were dehydrated, embedded, and sectioned at the UT Southwestern Electron Microscopy Core. TEM images were acquired on a JEOL 1400+ microscope using an AMT camera. G-ratios were calculated with AxonDeepSeg (*61*) and manually verified.

### Lentivirus production

The ss-RFP-GFP-KDEL plasmid (Addgene, Cat#128257) was cloned into a CMV promoter lentiviral backbone (Addgene, Cat#17448). Lenti-X 293T cells (Takara, Cat#632180) were co-transfected with the ss-RFP-GFP-KDEL lentiviral plasmid, psPAX2 (Addgene, Cat#12260), and pMD2.G (Addgene, Cat#12259) at a ratio of 2:1:1 using standard transfection protocols. After 6 hours, the transfection medium was replaced with DMEM supplemented with 1% FBS to collect lentivirus. Viral supernatants were collected at 48 and 72 hours post-transfection. The viral supernatants were pooled and mixed using Lenti-X Concentrator (Takara, Cat#631232) according to the manufacturer’s instructions. The mixture was centrifuged at 1,500 × g for 45 minutes, and the viral pellet was resuspended in DMEM-SATO medium to obtain a 40 × concentrated viral stock for subsequent infections.

### ER-phagy flux detection

OPCs were seeded onto 24-well glass-bottom plates (Cellvis, Cat# P24-1.5P). Cells were infected with 5 µL of concentrated lentivirus expressing ss-RFP-GFP-KDEL per well in oligodendrocyte differentiation medium. (DMEM-SATO supplemented with 4.2 μg/mL forskolin, 10 ng/mL CNTF, and 40 ng/mL T3). After 24 hours, the medium was replaced with fresh differentiation medium, and OPCs were allowed to differentiate for a total of 8 days. Live-cell imaging was performed using a Nikon CSU-W1 dual-camera spinning disk confocal microscope equipped with a 37°C chamber and 5% CO₂ supply. Cells displaying mature oligodendrocyte morphology with both red and green fluorescence signals were selected for imaging and analysis.

### Quantitative proteomics

Oligodendrocytes (∼5,000,000/dish) cultured in two 15-cm dishes were briefly rinsed with DPBS and lysed in 500 μl RIPA buffer supplemented with 1× cOmplete protease inhibitor cocktail and 1× PhosSTOP phosphatase inhibitors. Protein concentration was determined using a BCA assay, and 400 μg proteins were used for quantitative mass spectrometry (MS). The samples were processed at the UT Southwestern Proteomics Core. Protein samples were reduced with TCEP and alkylated with iodoacetamide, followed by overnight trypsin digestion using S-Trap microcolumns (Protifi). Peptides were dried, reconstituted in 100 mM TEAB, and labeled with TMT. Excess label was quenched with 5% hydroxylamine, and labeled peptides were pooled and fractionated into eight fractions using Pierce High pH Reversed-Phase spin columns (Thermo Fisher Scientific, Cat#84868). Fractions were dried and reconstituted in 2% acetonitrile with 0.1% TFA prior to LC-MS/MS analysis. Peptides were separated on a 75 μm × 75 cm EasySpray column using an Ultimate 3000 RSLC-Nano system coupled to an Orbitrap Fusion Lumos. A 180-min linear gradient from 0–28% buffer B (A: 2% acetonitrile, 0.1% formic acid; B: 80% acetonitrile, 10% trifluoroethanol, 0.1% formic acid) was used. MS data were acquired in positive ion mode with the 120,000 resolution for full scans, followed by top-speed SPS-MS3 acquisition. MS2 used CID at 35% collision energy, and MS3 used HCD at 55%; dynamic exclusion was 25 s. Data were analyzed with Proteome Discoverer 3 using Sequest HT against the UniProt mouse proteome. Search parameters included 10 ppm precursor and 0.6 Da fragment mass tolerance, up to three missed trypsin cleavages, carbamidomethylation of cysteines and TMT labeling of N-termini/lysines as fixed modifications, and methionine oxidation as variable. Peptide FDR was set at 1%. GO enrichment analysis was conducted with Gorilla (*62*), using a ranked protein list that was filtered for a >1.5-fold change in *Atg5* KO versus control, with a significance threshold of *p* < 0.05. Subcellular localization assignments for these differentially enriched proteins were manually annotated based on UniProt database entries.

### Lipidomics

Oligodendrocytes (∼5,000,000/dish) cultured in 15-cm dishes were harvested and pelleted. Prior to extraction, samples were spiked with isotopically labeled lipid standards (UltimateSPLASH ONE and SphingoSPLASH I, Avanti Polar Lipids). Lipids were extracted using a methyl tert-butyl ether (MTBE) protocol, and the organic phase was dried under nitrogen and reconstituted in 100 μL chloroform/methanol (1:1, v/v) for analysis. Protein content of the remaining pellet was determined using a BCA assay. Extraction controls included an Extraction Blank (no sample or standards) and an Extracted IS (standards only). LC-MS/MS was performed on an Acquity Premier UHPLC system (Waters) coupled to a QTRAP 6500 mass spectrometer (SCIEX). Lipids were separated via HILIC chromatography on a Luna NH2 column (Phenomenex) using mobile phases A (acetonitrile/water/hexane, 92:6:2 v/v/v, +10 mM ammonium acetate, pH 9.3) and B (acetonitrile/water, 1:1 v/v, +10 mM ammonium acetate, pH 9.3). Quality control included pooled QCs (aliquots from all samples) injected every five samples, IS QCs, and porcine sulfatides (1 μg/mL) to monitor retention time and signal stability. LC-MS/MS peak areas were integrated using MultiQuant Software v3.0.3 (SCIEX), normalized to internal standards and protein content, log_10 transformed, and auto-scaled. Lipids missing in ≥2/3 of a sample group were excluded from analysis.

### Cholesterol measurement

*In vitro* differentiated oligodendrocytes were collected and resuspended in PBS. The cell suspension was divided into two fractions. One fraction was mixed with an equal volume of chloroform:methanol (2:1, v/v) for lipid and cholesterol extraction. The other fraction was mixed with an equal volume of RIPA buffer for protein extraction and quantification. For lipid and cholesterol extraction, samples were centrifuged, and the lower organic phase was carefully transferred to a clean tube and air-dried overnight. Dried lipids were subsequently used for cholesterol quantification with the Amplex Red Cholesterol Assay Kit (Thermo Fisher Scientific, Cat#A12216) following the manufacturer’s instructions. Protein concentration in the RIPA fraction was measured with the BCA kit for normalization.

### Quantification and statistical analysis

All statistical analyses except for proteomic and lipidomic data (see Method Details) were performed using GraphPad Prism (v9 or v10) or RStudio. Data are presented as mean ± SEM. The sample size (“*n*”) for each experiment is indicated in the main text and corresponding figure legends. Comparisons between two groups were conducted using two-tailed Student’s *t*-tests. Western blot quantifications between two groups were analyzed using one-sample *t*-tests. The empirical distributions of two samples were compared using the Kolmogorov-Smirnov test. For analyses involving multiple groups, one-way or two-way ANOVA followed by Tukey’s *post hoc* multiple comparisons tests were applied. Statistical significance was defined as follows: \**p* < 0.05, \*\**p* < 0.01, \*\*\**p* < 0.001, and \*\*\*\**p* < 0.0001; ns, not significant.

